# *In vivo* regulation of fluorescent fusion proteins by engineered kinases

**DOI:** 10.1101/2021.03.26.433940

**Authors:** Katarzyna Lepeta, Chantal Roubinet, Oguz Kanca, Amanda Ochoa-Espinosa, Dimitri Bieli, Clemens Cabernard, Markus Affolter, Emmanuel Caussinus

## Abstract

Reversible protein phosphorylation by kinases in extensively used to control a plethora of processes essential for proper development and homeostasis of multicellular organisms. One main obstacle in studying the role of a defined kinase-substrate interaction is that kinases form complex signaling networks and most often phosphorylate multiple substrates involved in various cellular processes. In recent years, several new approaches have been developed to control the activity of a given kinase. However, most of them fail to regulate a single protein target, likely hiding the effect of a unique kinase-substrate by pleiotropic effects. To overcome this limitation, we have created protein binder-based engineered kinases for direct, robust and tissue-specific phosphorylation of target fluorescent protein fusions *in vivo*. We show that synthetic Rok kinases, based on the *Drosophila* ortholog of Rho-associated protein kinase (ROCK), are functional enzymes and can activate myosin II through phosphorylation of Sqh::GFP or Sqh::mCherry in different morphogenetic processes in a developing fly embryo. We next use the system to study the impact of actomyosin activation specifically in the developing tracheal branches and showed that ectopic activation of actomyosin with engineered Rok kinase did not prevent cell intercalation nor the formation of autocellular junctions. We assume that this approach can be adapted to other kinases and targets in various eukaryotic genetic systems.

## Introduction

Reversible protein phosphorylation by kinases constitutes one of the most common regulatory mechanisms used to control cell signaling across eukaryotes. There are about 500 mammalian and about 250 *Drosophila* protein kinases encoded in the respective genomes^1,2^. Eukaryotic protein kinases catalyze the transfer of a phosphate group from ATP to the substrate protein to control protein activity and localization, providing information transfer in the cell in response to an appropriate signal^3^. Consequently, protein kinases orchestrate multiple intracellular processes such as cell determination and differentiation, cell migration and response to environmental cues or stress, allowing for proper development and homeostasis of multicellular animals. Dysregulation of kinase signaling is a hallmark of numerous diseases including cancer^4,5^. Thus, synthetic approaches for regulating kinase activity, be it for dissecting cell signaling pathways, for building synthetic biology modules, or for therapeutic purposes, are of great need^6^.

According to their substrate specificity, eukaryotic protein kinases are generally subdivided into three groups, serine/threonine kinases, tyrosine kinases and dual-specificity kinases^1,7,8^. One main obstacle in studying the role of a defined kinase-substrate interaction is that kinases form complex signaling networks and most often phosphorylate multiple substrates involved in various cellular processes. Hence, regulating the phosphorylation of a single protein target and dissecting the resulting phenotype is both a very desirable and a very challenging task. In recent years, several new approaches for manipulating kinase activity or localization have been developed. Those include small molecule kinase inhibitors, engineered isolated kinase domains or light-controlled regulation of kinase activity (extensively reviewed in^6,9–11^). However, although some of these approaches allow for precise temporal and spatial control of the activity of a given kinase (opto-based tools), most of them fail to regulate a single protein target, likely hiding the effect of a unique kinase-substrate by pleiotropic effects. In addition, optical manipulation is only possible up to a limited tissue depth precluding the use of opto-based in non-light permeable tissues and in many cases require the use of UV light (the vast majority of photolabile caging moieties introduced into kinases/small molecule kinase inhibitors), hindering its extensive use *in vivo* due to the toxicity and limited tissue penetration of UV light ^9,12,13^. Photoswitchable kinases (generated by inclusion of an allosteric photosensory domain or a photodissociable dimeric domain into the kinase of interest) employ longer wavelengths (400–500 nm) for isomerization compared to the UV light used for the uncaging reactions; however, to date, only a limited alteration of kinase activity between the active and inactive form has been achieved, limiting efficient control over kinase activity.

Protein binders, derived from either single chain antibodies, so-called nanobodies (reviewed in ^14,15^), designed ankyrin repeat proteins (DARPins)^16–18^, or from other scaffolds ^19,20^, can be used to direct enzymes to specific and unique substrates through direct protein-protein interaction. In the last decade, multiple small, genetically encodable protein binders against fluorescent proteins have been isolated and functionalized by fusing them to various effector domains, providing a growing toolkit to study protein function in a more direct manner in living organisms (extensively reviewed in ^11,21,22^).

Here, we characterize protein binder-based engineered kinases for direct, robust and tissue-specific phosphorylation of target fluorescent protein fusions *in vivo* (Fig. 1a). For these series of experiments, we use Rok kinase, the *Drosophila* ortholog of Rho-associated protein kinase (ROCK)^23,24^, which is essential for myosin II activation. Rok is a serine/threonine (S/T) kinase regulated by the small GTPase Rho1^25^. In its inactive state, the C-terminal Pleckstrin homology (PH) domain of Rok inhibits the activity of the N-terminal kinase domain (N-Rok). Upon binding of the activated form of Rho1, Rok changes its conformation and N-Rok becomes active^26^. Rok acts in synergy with myosin light-chain kinase (MLCK) to activate Spaghetti squash (Sqh/Myosin II Regulatory Light Chain) with a sequential phosphorylation on Ser21 and Thr20, which in turn activates myosin II^23,24,27^ (Fig. 1b). Both Rok and MLCK are antagonized by Flapwing (Flw), a serine/threonine protein phosphatase that can inactivate Sqh^28^. Myosin II is the major contractile protein complex of non-muscle cells, and as such a prominent target of Rok/ROCK. The other known substrates of Rok are Myosin binding subunit, Diaphanous/Diaphanous-related formin, α-adducin, LIM-kinase1, MAP2, Tau and Combover^27,29–32^.

**Figure 1.**
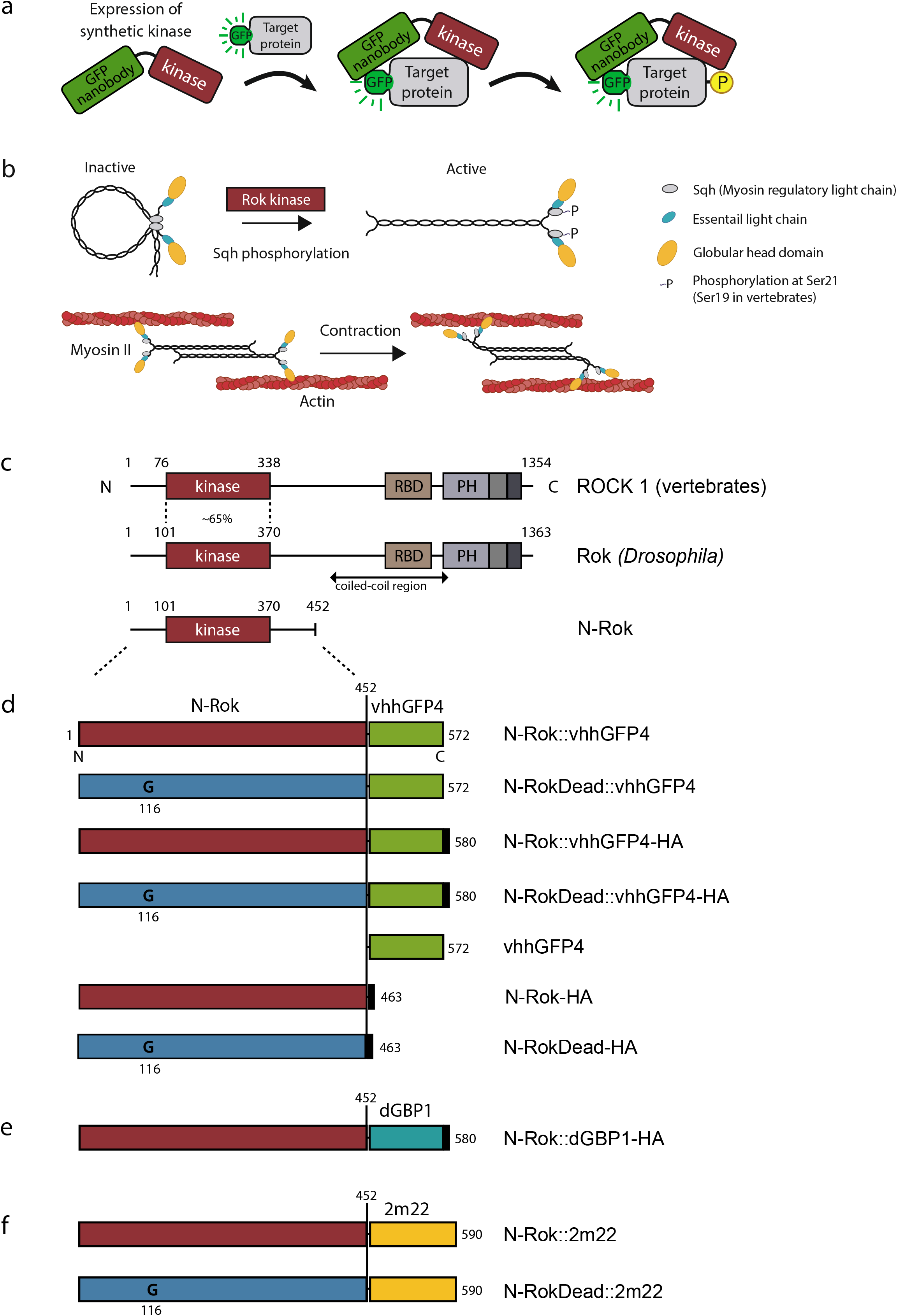
Schematic illustration of engineered synthetic kinases. (a) Overview of engineered kinase work concept. A synthetic kinase uses a small fluorescent protein binder (here: vhhGFP4, a GFP nanobody) to bring a constitutively active kinase domain (kinase) in close proximity to a fluorescent fusion portein (target). The persistence of the kinase domain around the fluorescent fusion protein allows for an efficient phosphorylation (P) of the target. (b) Schematic illustration of non-muscle myosin II activation and actomyosin contractility. Non-phosphorylated Sqh/Myosin-II regulatory light chain (MRLC) assembles into inactive compact molecule through a head to tail interaction. Reversible phosphorylation of Sqh at Ser21 (Ser19 in mammalian MRLC) results in myosin II molecule unfolding, allowing association with other myosin II molecules in an antiparallel fashion and binding to the actin filaments through the head domains. An ATP-dependent conformational change in myosin II drives the actin filament sliding in an anti-parallel manner and results in contraction. (c) The structure of mammalian and *Drosophila* ROCK proteins. The *Drosophila* Rok kinase region shares ~65% identity with the corresponding isolated domain of mammalian ROCK1^27^. The N-terminal kinase region (N-Rok) of Drosophila Rok (amino acid 1 to 452) was used for synthetic kinase constructs shown in (d-f). (d) Linear representation of N-Rok::vhhGFP4, N-RokDead::vhhGFP4, N-Rok::vhhGFP4-HA, N-RokDead::vhhGFP4-HA, vhhGFP4, N-Rok-HA and N-RokDead-HA. (e) Linear representation of N-Rok::dGBP1, in which a destabilized GFP-binding nanobody (dGBP1)^33^ substitutes vhhGFP4 from d. (f) Linear representation of N-Rok::2m22, N-RokDead::2m22 in which a DARPin recognizing mCherry substitutes vhhGFP4 from d. In some constructs human influenza hemagglutinin tag (HA; black squares) was added to the C-terminus of the synthetic kinases to allow detection by immunofluorescence. N-RokDead contains a K116G single amino acid substitution (G) that abolishes catalytic activity. The proteins are aligned vertically with the N-Rok domain. Numbers refer to amino acid positions from N-terminus (N) to C-terminus (C).

In order to generate an engineered Rok kinase that allows for targeted fluorescent fusion protein phosphorylation with high specificity, we fused the catalytic domain of Rok kinase (Fig. 1c) with a nanobody either directed against GFP and its close derivatives (vhhGFP4, Fig. 1d)^16^, a destabilized GFP-binding nanobody (dGBP1, Fig. 1e)^33^ or a DARPin against mCherry (2m22, Fig. 1f)^17^. We show that synthetic Rok kinases are functional enzymes and can activate myosin II through phosphorylation of Sqh::GFP or Sqh::mCherry in different morphogenetic processes in a developing fly embryo, namely in dorsal closure and during the development of the *Drosophila* tracheal system. We show that tissue-specific expression of the engineered Rok kinase together with Sqh::GFP leads to the formation of ectopic clusters of activated actomyosin (and ectopic tension forces) around the Sqh foci, leading to perturbed cell migration and failure in proper embryo closure and tracheal branching.

Lastly, we used the Rok kinase fused with the destabilized GFP-binding nanobody (dGBP1) to address a long-standing question regarding the need for actomyosin regulation to allow for tracheal cell intercalation. In sharp contrast to the actomyosin-driven junctional rearrangements observed in many intercalation events in drosophila morphogenesis^34–43^, tracheal cell intercalation occurs passively and does not require actomyosin contractility^44^. Excessive phosphorylation of Sqh::GFP by synthetic N-Rok::dGBP1^ZH-86Fb^ in the *kni* domain did not result in a general failure of stalk cells intercalation, despite evident clusters with ectopic points of tension in the areas most likely corresponding to cell junctions; stalk cell intercalation and autocellular junction formation were seen in most branches upon actomyosin activation. However, ectopic activation of actomyosin caused occasional aberrant contacts establishment or – in more severe cases – led to detachment of parts of the cells from the rest of the branch, resulting in an aberrant branching network. Our results provide evidence that tip cell migration, dorsal branch extension and cell intercalation in most cases can occur despite excessive actomyosin activation, supporting the notion that tip-cell pulling force is a key player in proper formation of the dorsal branch and that this force is able to overcome some counteracting excessive points of tension.

## Materials & Methods

### Generation of engineered kinase constructs

To generate pUASTattB_N-Rok::vhhGFP4 and pUAST_N-Rok::vhhGFP4, the N-terminal kinase region of Drosophila Rok (N-Rok) was fused to vhhGFP4 and inserted into respectively pUASTattB vector^45^ or pUAST with EcoRI and XbaI. In pUASTattB_N-Rok::vhhGFP4-HA, a C-terminal Human influenza hemagglutinin tag (HA) was added. In N-RokDead variants, a K116G single amino acid substitution (K->G) that abolishes catalytic activity^24^ was introduced by site-directed mutagenesis. In pUASTattB_N-Rok::dGBP1, vhhGFP4 was replaced by a destabilized GFP-binding nanobody (dGBP1)^33^. To generate pUASTattB_N-Rok::2m22 and pUASTattB_N-RokDead::2m22, the N-Rok variants were fused to a DARPin recognizing mCherry (2m22)^17^. See Supplementary Data 1 for sequences.

### Immunofluorescence staining of S2 cells

Stable *Drosophila* S2 cell line expressing Sqh::GFP (S2 Sqh::GFP)^46^ was grown in Schneider’s *Drosophila* medium (Invitrogen) supplemented with 10% heat-inactivated Fetal Bovine Serum (FBS) and Penicillin-Streptomycin (at a final concentration of 50 units penicillin G and 50 μg streptomycin per ml of medium) at 25°C. Cells were transfected with indicated plasmids using FuGENE HD (Roche) according to the manufacturer’s instructions. Cells were fixed with 4% PFA for 30 min at RT 40 hours after transfection and processed for immunofluorescence using standard procedures. Briefly, blocking and permeabilization was performed in TBS (20 mM Tris-HCl, pH 7.5, 154 mM NaCl, 2 mM EGTA, 2 mM MgCl2) containing 2% BSA and 0.02% saponin for 1.5 hour. Staining was done using the same solution. Antibodies used were mouse anti-HA (Roche; 12CA5, 1:200), rat antitubulin (1:200), rabbit anti-Sqh2P (Thermo Scientific, 1:200), rat anti-Sqh2P (1:200)^47^, rabbit anti-Sqh1P (1:50)^47^. Cells were resuspended in Vectashield mounting medium containing DAPI, transferred to microscope slides and covered with 22 mm^2^ coverslip. Statistical significance was determined with Student’s t test using GraphPad Prism software. The Prism convention is used: n.s. (P > 0.05), *(P ≤ 0.05), **(P ≤ 0.01), ***(P ≤ 0.001) and ****(P ≤ 0.0001).

### Experimental model

*Drosophila melanogaster* was cultured using standard techniques at 22-25°C on standard corn meal *Drosophila* medium supplemented with yeast. For virgin collection fly vials were kept at 18°C for the period of collecting; both male and female animals were used.

### Generation of transgenic fly lines

To achieve the same level of expression of the different engineered kinase constructs with the same Gal4 driver, we used the phiC31 integrase^45^ system and a unique site of integration (ZH-86Fb)^45^ on third chromosome to generate transgenic lines carrying *N-Rok::vhhGFP4*^ZH-86Fb^, *N-RokDead::vhhGFP4*^ZH-86Fb^, *N-Rok::vhhGFP4-HA*^ZH-86Fb^, *N-RokDead::vhhGFP4-HA*^ZH-86Fb^, *N-Rok-HA*^ZH-86Fb^, *N-RokDead-HA*^ZH-86Fb^, *N-Rok::2m22*^ZH-86Fb^, *N-RokDead::2m22*^ZH-86Fb^ and *N-Rok::dGBP1*^ZH-86Fb^. The *N-Rok::vhhGFP4*^Vi^ fly line was generated by random genomic insertion of N-Rok::vhhGFP4 by P element mediated transformation^48^.

### Drosophila stocks and genetics

All fly strains used in this study are listed in the Resources Table.

For tissue-specific Sqh::GFP phosphorylation during dorsal closure females harboring *UAS*-indicated engineered kinase construct (all listed in the Resources Table) were crossed with *sqh_Sqh::GFP enGal4 UAS_mCherry-nls* or with *enGal4 sqh_Sqh::mCherry* males; or with *enGal4 UAS_mCherry-nls/ (CyO DfdGMR_YFP)* males (control without the target). For the controls without the engineered kinases *sqh_Sqh::GFP enGal4 UAS_mCherry-nls/+* or *sqh_Sqh::GFP enGal4 UAS_mCherry-nls/P{2xTb[1]-RFP}CyO* (from Bloomington stock #36336) embryos were used. For evaluation of engineered Rok kinase in trachea, *sqh[AX3]; {sqh_Sqh::GFP} {btl-Gal4} {UAS-mCherry-nls}* females were crossed with males expressing *UAS-*indicated engineered kinase variants.

To evaluate engineered Rok activity on other targets than Sqh, following flies were used: *Tkv::YFP* females (or yw for control) were crossed with *enGal4 UAS_RhoK::vhhGFP4*^Vi^/*(CyO DfdGMR_YFP)* or with *enGal4 UAS_RhoKDead::vhhGFP4/TSTL* males; *enGal4 UAS_RhoK::vhhGFP4*^Vi^/*(CyO DfdGMR_YFP)* (or yw for control) females were crossed with shg::GFP males.

### Antibody staining of Drosophila embryos

Embryos were collected on grape juice agar plates supplemented with yeast paste after overnight egg laying at 25°C and processed for immunofluorescence using standard procedures. Briefly, embryos were collected on a nitex basket, dechorionated in 50% bleach (Sodium hypochlorite, stock solution 13 % w/v technical grade, AppliChem GmbH), washed thoroughly with dH_2_O and fixed in 50:50 heptane: 4% paraformaldehyde solution for ~20 minutes with vigorous shaking. Embryos were devitellinized by replacement of the fix phase with the same volume of MeOH and vigorous hand shaking for 1 minute. After washing 4 times with MeOH, the embryos were either stored at −20°C to accumulate enough material or directly subjected to staining. After brief re-hydration of fixed embryos in 1:1 1xPBS:MeOH solution, embryos were rinsed with 1xPBS, washed 4 x 15 min in PBST (PBS + 0.3% Triton X-100), blocked in PBTN (PBST + 2% normal goat serum) for 1h and incubated overnight with primary antibodies in PBTN at 4°C on rotator. Embryos were rinsed twice and then washed 4x 20 min with PBST and incubated with secondary antibodies in PBTN for 2-3 hours, washed as before and rinsed 2x with PBS. Embryos were mounted on microscope slide in Vectashield mounting medium (H-1000, Vector Laboratories) and covered with 22 mm^2^ coverslip.

The monophosphorylation on serine 21 in embryos was detected with a guinea pig anti-Sqh1P antibody serum in 50% glycerol (gift from R. Ward) at 1:400 concentration or a rabbit anti-phospho-Myosin Light Chain 2 (Ser19) (#3671, Cell Signaling Technology) at 1:50 concentration. DCAD2 rat at 1:25 (DSHB), vermiform rb 1:300 (^49^). Secondary antibodies used were Alexa Fluor 488/Fluor 561/Fluor 647 coupled (Thermo Fisher Scientific, 1:500).

### Embryo mounting for live time-lapse imaging

Embryos were dechorionated in 30-50% bleach (Sodium hypochlorite, Sodium hypochlorite, stock solution 13 % w/v technical grade, AppliChem GmbH) and extensively rinsed in water. Embryos at desired stages were selected manually on grape juice agar plate under a dissection scope and mounted either with heptane-glue method (I) or on glass-bottom dish (II).

(I) Briefly, selected embryos were manually aligned on agar plates and attached to heptane-glue coated coverslips by gently pressing the trail of glue against the embryos. A piece of permeable bio-foil (bioFOLIE 25, In Vitro System and Services, GmbH) was stretched over a custom-made metal slides with a hole, and embryos were mounted in a drop of halocarbon oil 27 (Sigma) between the foil and the coverslip (see^50^ for details of the procedure).

(II) Embryos were transferred with cleaned forceps to a glass-bottom dish (MatTek, 35 mm dish, co. 1.5 coverslip, uncoated, P35G-1.5-10-C). After adding 1xPBS, embryos were gently rolled to the desired positions using a cut gel loading pipette tip (flexible, to minimize embryo disruption). Properly dechorionated embryos adhere to the glass-bottom; the part to be imaged was facing down.

### Confocal and time-lapse imaging

Images of fixed embryos were acquired with 1024×1024 frame size with a 40x / 1.25NA oil objective (HCX PL APO) on a Leica TCS SP5 II inverted scanning confocal microscope with 2.5 zoom. A series of z-stacks was acquired for each embryo at 0.5 μm step size using a 488 nm, 561 nm and 633 nm laser line.

Time-lapse sequences of dorsal closure or whole trachea development were imaged under either a) Point Scanning Confocal Zeiss LSM880 AiryScan inverted microscope (Confocal mode) with PLAN APO 40x / 1.2NA objective (LD LCI PLAN APO, Imm Corr DIC M27, water immersion) with 0.9 zoom using Zen Black software, at 25°C or b) Leica TCS SP5 II inverted scanning confocal microscope with 40x / 1.25NA oil objective (HCX PL APO) (embryos mounted in oil with heptane glue method) or 40x / 1.10NA water objective (HCX PLAN APO CS) (glass-bottom dish mounted embryos) using LAS AF software. A series of z-stacks was acquired for each embryo at 0.5 μm - 1μm steps using a 488 nm and 561 nm laser line. Imaging was carried out at 5 min intervals (movies corresponding to Fig. 3b, c, Fig. 6 & Fig. 7) or 10 min intervals (movies corresponding to Fig. 3a, Fig. 4b, e & Fig. 5b).

**Figure 2.**
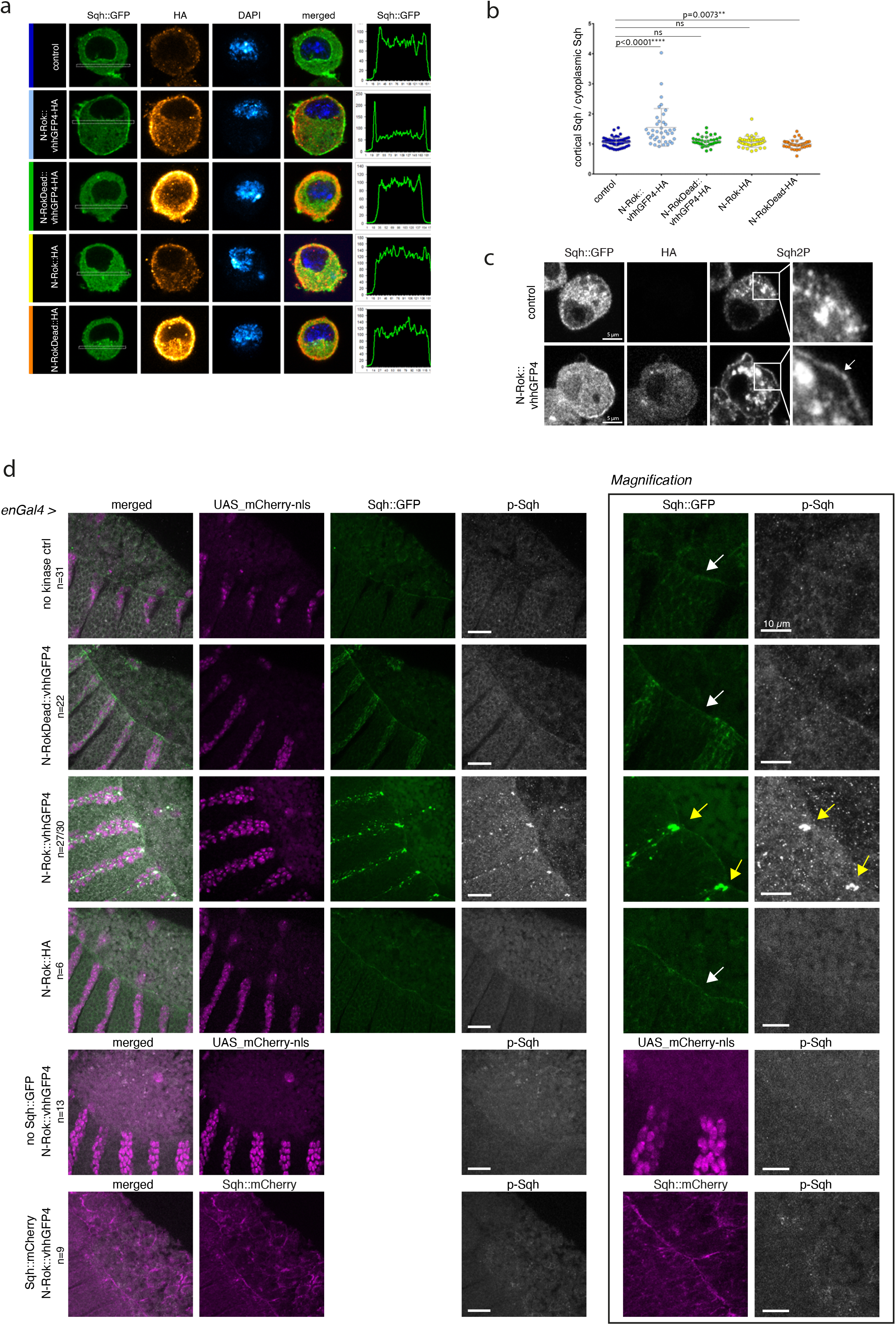
N-Rok::vhhGFP4 is a functional enzyme and efficiently phosphorylates Sqh::GFP *in vivo* in a tissue specific manner. (a) Expression of N-Rok::vhhGFP4 efficiently recruits Sqh::GFP at the cell cortex in interphasic cells. Stable S2 cell line expressing Sqh::GFP, transfected with N-Rok::vhhGFP4-HA, N-RokDead::vhhGFP4-HA, N-Rok-HA or N-RokDead-HA. Sqh::GFP shown in green, anti-HA staining in red (transfected cells), anti-tubulin staining in blue (discrimination of interphasic from mitotic cells), DAPI in dark blue. Graphs on the right show the ratio of cortical Sqh::GFP/cytoplasmic Sqh::GFP of a representative cell shown for each genotype. (b) Quantification of the data shown in a. n=218 cells, data from 3 independent experiments. (c) Stable S2 cell line expressing Sqh::GFP transfected with N-Rok::vhhGFP4-HA and stained with anti-Sqh2P antibody. The right panel shows magnification of the indicated cell region, white arrow points to the enrichment of phospho-Sqh signal at the cell cortex. (e) Panels show lateral views of fixed *Drosophila* embryos at stage 13-14 (dorsal closure) expressing Sqh::GFP and the synthetic kinase in the engrailed domain (visualized by co-expression of mCherry-nls). Embryos were stained with anti-phospho-myosin antibody. The right panel shows magnification of the respective Sqh::GFP and p-Sqh images for each of the embryo genotypes shown on the left. White arrows point the actomyosin cable around the dorsal hole, yellow arrows point to Sqh::GFP and p-Sqh foci. For every expressed synthetic kinase, the number of considered embryos is indicated (n). For N-Rok::vhhGFP4 the n number is given as a proportion of embryos in which clear p-Sqh clumps corresponding to Sqh::GFP foci were clearly visible to the total number of embryos included in the analysis. Scale bar: 20 μm, in the zoomed panel – 10 μm.

**Figure 3.**
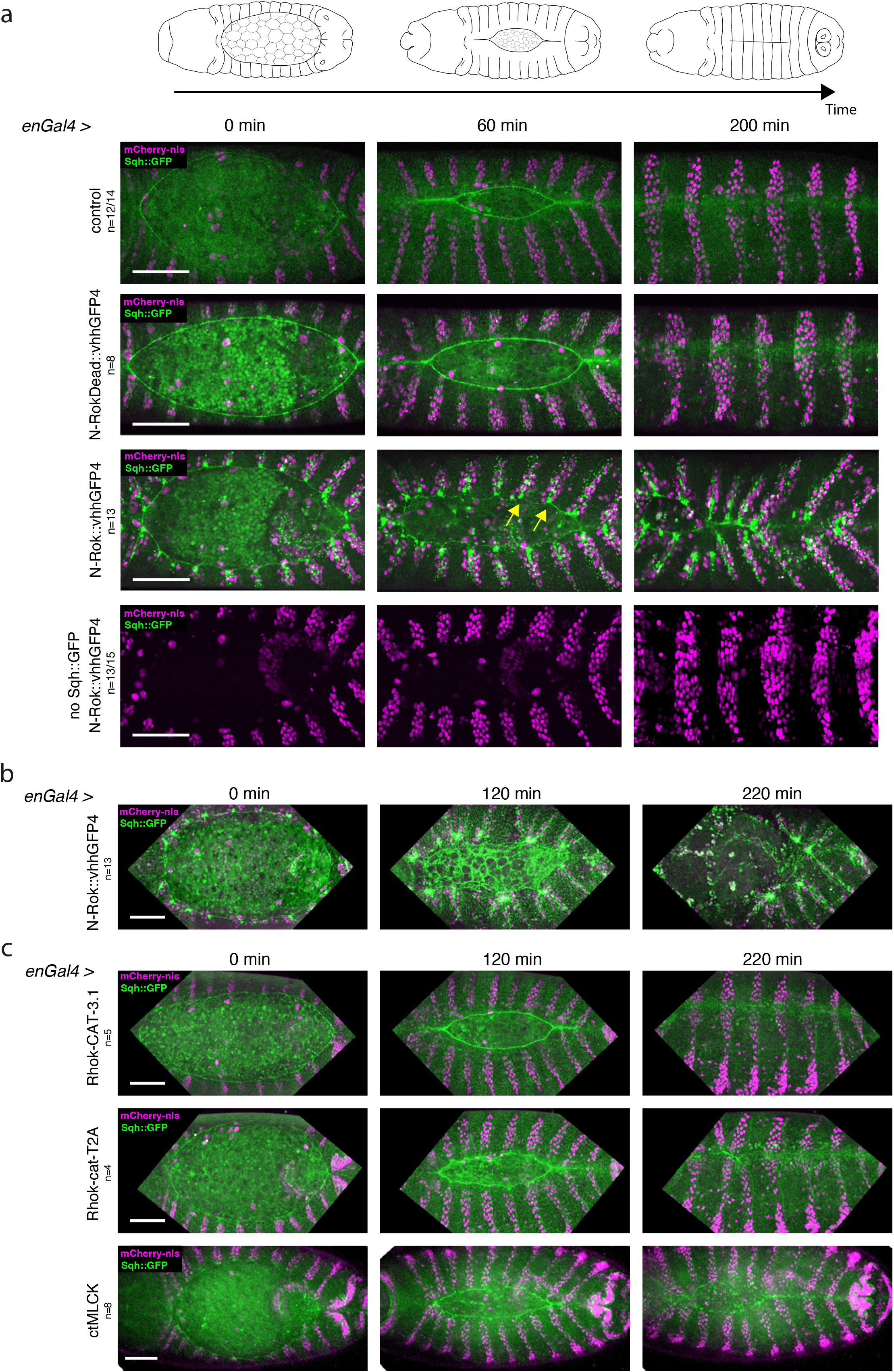
Synthetic Rok kinase modulates mechanical properties of cells through the phosphorylation of Sqh::GFP and myosin II activity. (a) Schematic illustration of dorsal closure process in the developing fly embryo. Dorsal closure was used as a model to assess myosin II activation by means of Sqh::GFP phosphorylation with synthetic Rok. In *sqh_Sqh::GFP* embryos, expression of N-Rok::vhhGFP4 leads to abnormal dorsal closure. All panels show stills from live-imaging with dorsal views of the developing *Drosophila* embryos at stages 13/14-16 (dorsal closure) expressing Sqh::GFP and variants of the synthetic kinase in the engrailed domain (visualized by co-expression of mCherry-nls). Note the yellow arrows pointing to the Sqh::GFP foci and actomyosin cable invaginations in N-Rok::vhhGFP4 panel. (b) Stills from live-imaging with dorsal views of the developing *sqh_Sqh::GFP* embryo expressing N-Rok::vhhGFP4 in the engrailed domain mounted with gluing to the coverslip technique to show more severe dorsal open phenotype than embryos imaged on a glass-bottom dish shown in (a). (c) Dorsal closure was used to compare N-Rok::vhhGFP4 with the previously published effectors that are known to activate myosin II. All panels show stills from live-imaging with dorsal views of the developing *sqh_Sqh::GFP* embryos at stages 13/14-16 (dorsal closure) expressing myosin II activating tool in the engrailed domain (visualized by co-expression of mCherry-nls). Only for Fig. 3b, c and corresponding movies, the embryos were imaged with gluing to the coverslip technique; all the other presented embryos were imaged on a glass-bottom dish. Scale bars: 50 μm. Images are representative of n embryos indicated for every expressed synthetic kinase variant.

**Figure 4.**
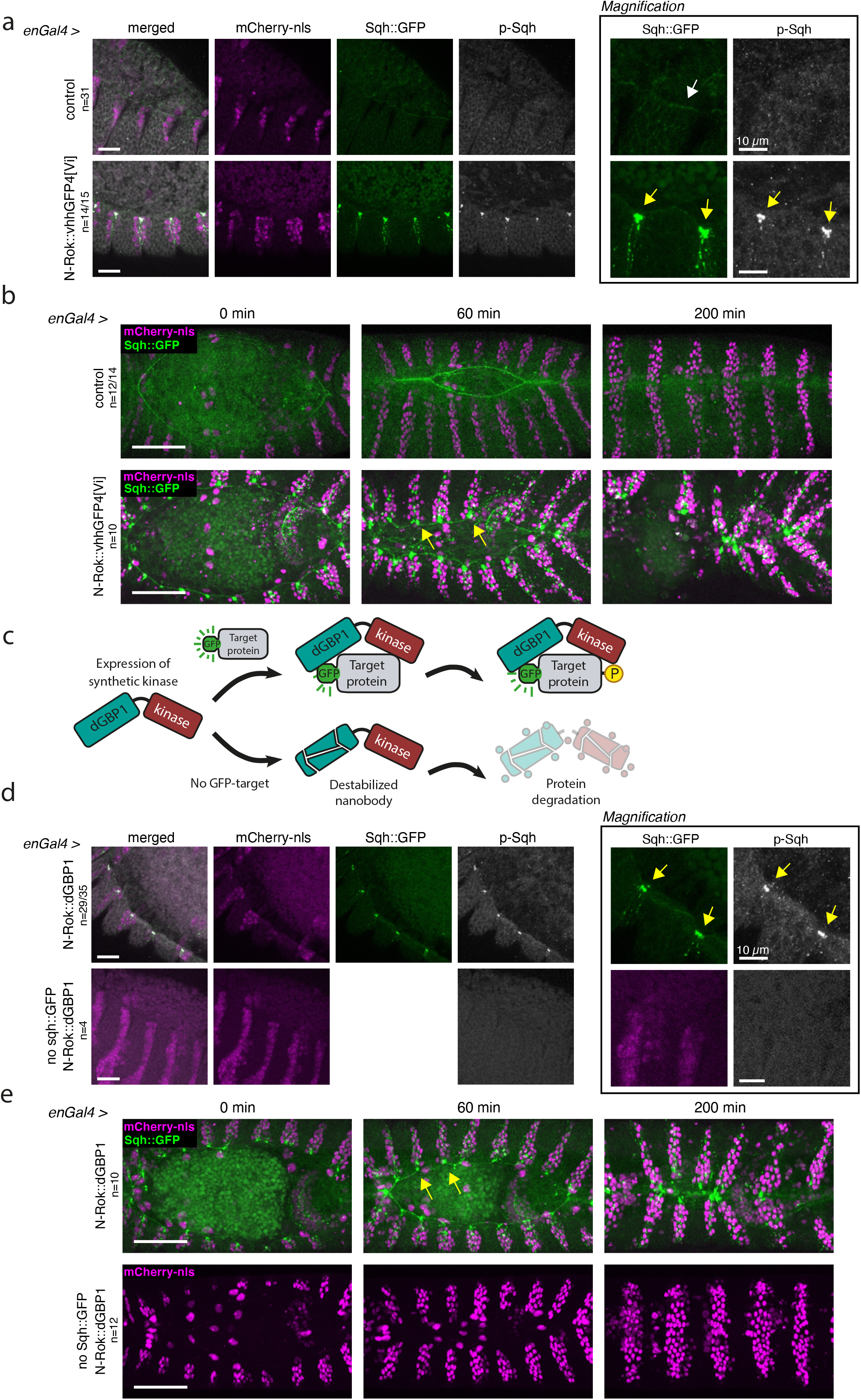
N-Rok::vhhGFP4^Vi^ and N-Rok::dGBP1 are optimized variants of synthetic N-Rok. (a) Panels show lateral views of fixed *sqh_Sqh::GFP* embryos at stage 13-14 (dorsal closure) expressing N-Rok::vhhGFP4^Vi^ in the engrailed domain (visualized by co-expression of mCherry-nls). Embryos were stained with anti-phospho-myosin antibody. The panel on the right shows magnification of the respective Sqh::GFP and p-Sqh images for each of the presented embryos. White arrows point the actomyosin cable around the dorsal hole, yellow arrows point to Sqh::GFP and p-Sqh foci. b) Dorsal closure was used to compare functionality of N-Rok::vhhGFP4^Vi^ to N-Rok::vhhGFP4 (Fig. 3a). Panels show stills from live-imaging with dorsal views of the developing *sqh_Sqh::GFP* embryos at stages 13/14-16 (dorsal closure) expressing N-Rok::vhhGFP4^Vi^ in the *engrailed* domain (visualized by coexpression of mCherry-nls). Please note, that both in a and b the “control” images are the same as on Fig. 2e and 3a, respectively, to have a single reference image for all N-Rok variants used. (c) Schematic illustration of the synthetic kinase with destabilized GFP nanobody (dGBP1) work concept. In a similar way as on Fig. 1a, a synthetic kinase uses GFP to bring a constitutively active kinase domain (kinase) in close proximity to a fluorescent fusion substrate protein (target). The persistence of the kinase domain around the fluorescent fusion protein achieves an efficient phosphorylation (P) of the target. In the absence of the GFP-target, dGBP1 nanobody is destabilized, and the whole nanobody-kinase fusion protein is targeted to degradation. (d) Panels show lateral views of stage 13-14 (dorsal closure) fixed *sqh_Sqh::GFP* or control yw embryos expressing N-Rok::dGBP1 in the engrailed domain (visualized by co-expression of mCherry-nls). Embryos were stained with anti-phospho-myosin antibody. The panel on the right shows magnification of the respective Sqh::GFP and p-Sqh images. Yellow arrows point to Sqh::GFP and p-Sqh foci. e) Dorsal closure was used to compare functionality of N-Rok::dGBP1 to N-Rok::vhhGFP4 (Figure 3a) and N-Rok::vhhGFP4^Vi^ (b). Panels show stills from live-imaging with dorsal views of the developing *sqh_Sqh::GFP* or control yw embryos expressing N-Rok::dGBP1 in the *engrailed* domain (visualized by co-expression of mCherry-nls). For every embryo genotype, the number of analyzed embryos is indicated (n). For N-Rok::dGBP1 the n number is given as a proportion of embryos in which clear p-Sqh clumps corresponding to Sqh::GFP foci were visible to the total number of embryos included in the analysis. Scale bar: 50 μm in b, e; 20 μm in a, d, in the zoomed panel – 10 μm.

**Figure 5.**
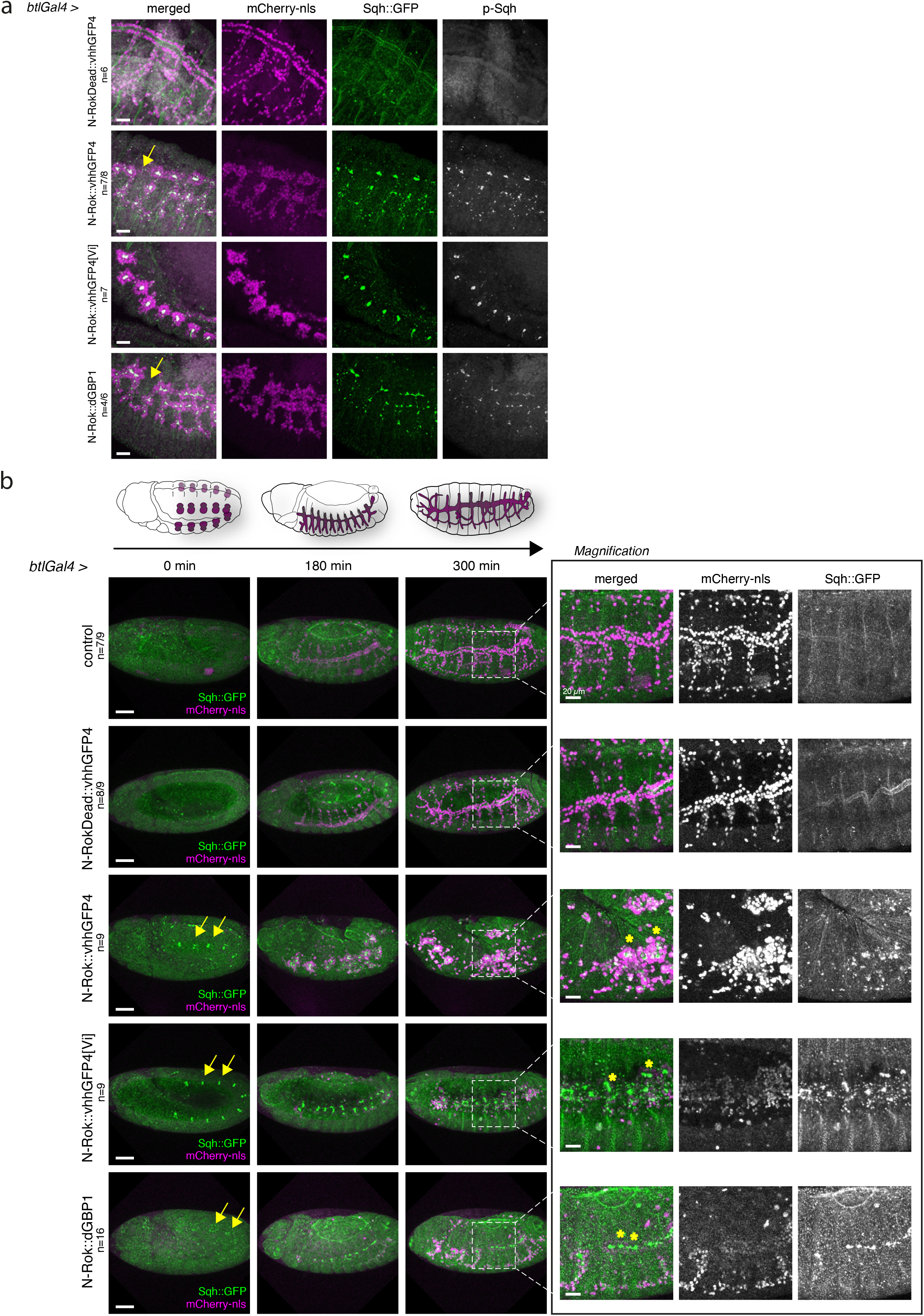
Tracheal system of Drosophila embryo develops abnormally due to the excessive myosin II phosphorylation by engineered N-Rok kinase. a) All panels show lateral views of fixed *sqh_Sqh::GFP* embryos at stage 14-16 expressing the indicated variant of synthetic Rok kinase in the tracheal system (*btl* expression domain visualized by coexpression of mCherry-nls). Embryos were stained with anti-phospho-myosin antibody. For every expressed synthetic kinase, the number of embryos analyzed is indicated (n). For N-Rok::dGBP1, the n number is given as a proportion of embryos in which clear p-Sqh clumps corresponding to Sqh::GFP foci were visible, to the total number of embryos included in the analysis. Scale bar: 20 μm. b) Aberrant tracheal development in *sqh_Sqh::GFP* embryos expressing the indicated variant of synthetic Rok kinase in the tracheal system. All panels show stills from live-imaging with lateral views of *sqh_Sqh::GFP* embryos at stages 11/12, 14 and 17, corresponding to the stages depicted on the schematic drawing above. The panel on the right shows magnification of the indicated region from the last stage. Yellow arrows point to Sqh::GFP foci visible already at stage 11/12. Scale bars: 50 μm, in the zoomed panel – 20 μm.

**Fig. 6.**
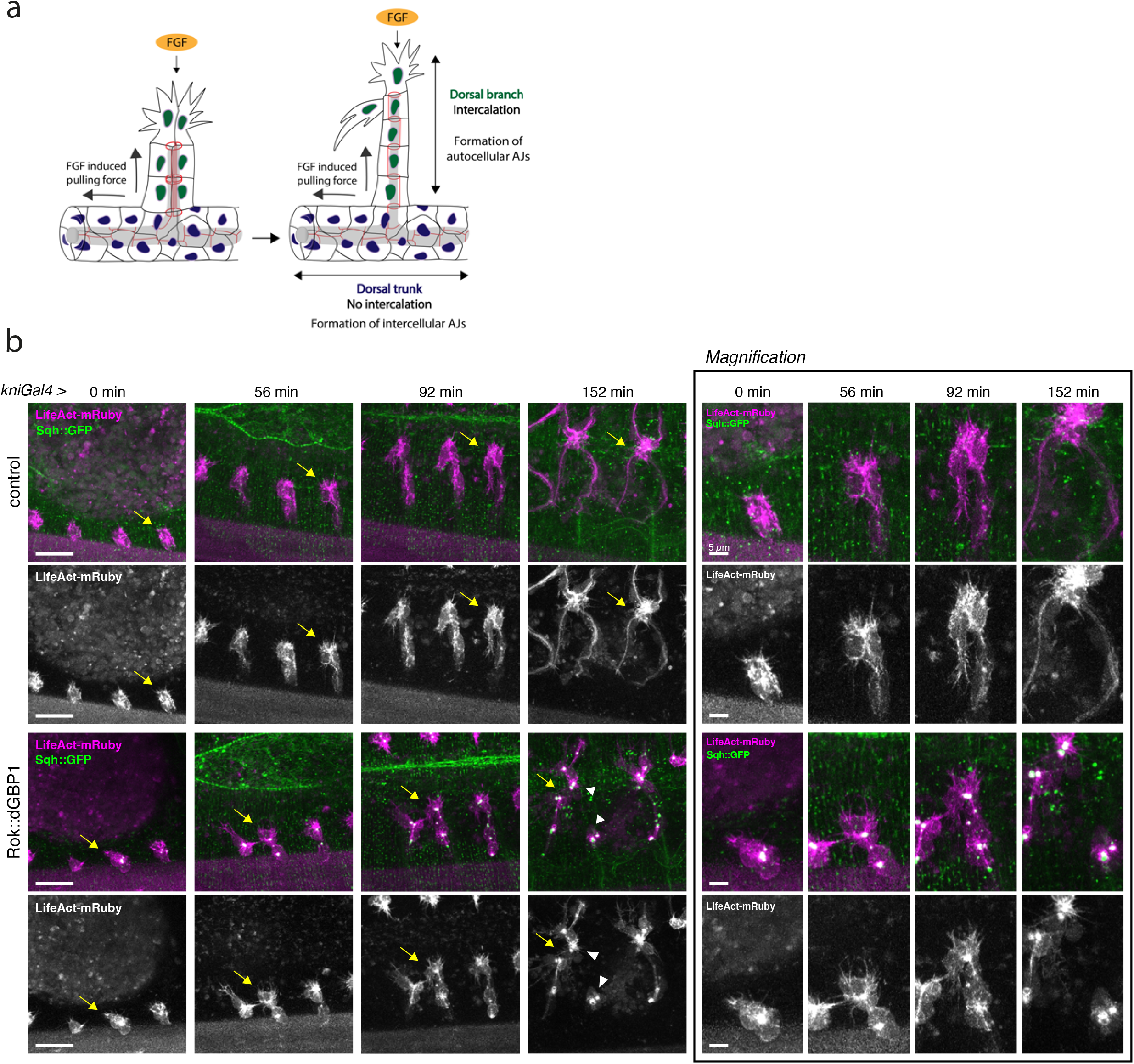
Effect of constitutive activation of actomyosin with N-Rok::dGBP1 in dorsal branches on filopodia formation and cell migration. (a) Schematic illustration of dorsal branch formation, showing cell junction rearrangements during cell intercalation. Cell intercalation is brought about by migration of tip cells towards the fibroblast growth factor (FGF) source. (b) All panels show stills from live-imaging with dorsolateral views of dorsal branches of *sqh_Sqh::GFP kniGal4* control embryos and *sqh_Sqh::GFP kniGal4 Rok::dGBP1* embryos; LifeAct-mRuby (red) was used to visualize pools of F-actin. Arrows indicate the same dorsal branch for all timepoints; arrowheads point to the tip cell detached from the stalks cells. Scale bars: 20 μm or 5 μm on the magnification panel.

**Fig. 7.**
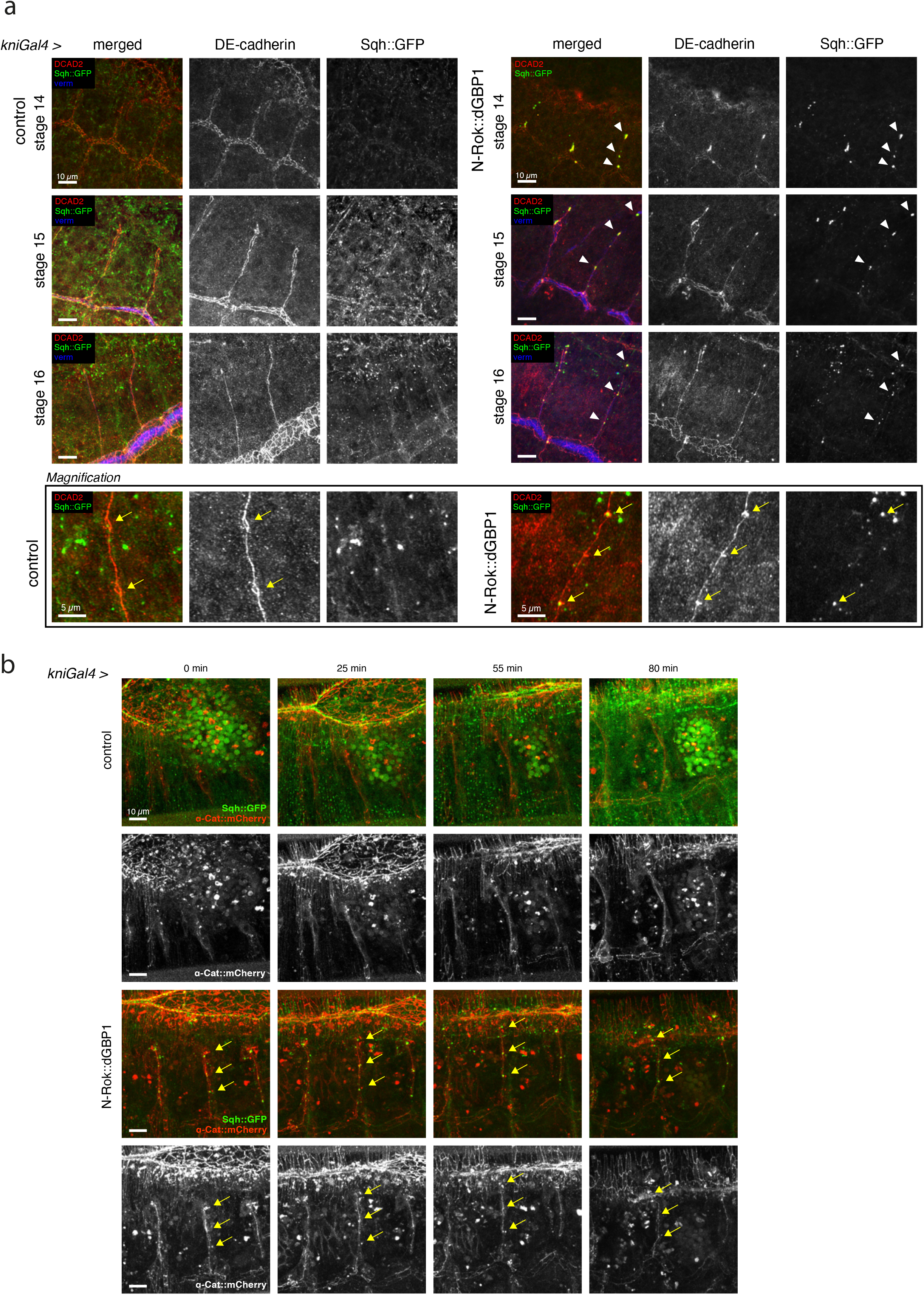
Effect of constitutive activation of actomyosin with N-Rok::dGBP1 in the *kni* domain on cell intercalation. (a) Confocal projections showing lateral views of fixed *sqh_Sqh::GFP kniGal4* control and *sqh_Sqh::GFP kniGal4 Rok::dGBP1* embryos stained for DEcadherin (red) and vermiform (blue). Note the presence of ectopic Sqh::GFP foci in the embryos expressing the engineered kinase (arrowheads). Bottom panel shows magnification of a single branch from the respective image of stage 16 embryos to the show characteristic pattern of lines and small rings of AJs, corresponding to auto- and intercellular AJs, respectively. Yellow arrows point to the rings corresponding to intercellular AJs. (b) Stills from time-lapse movies showing projections of dorsal branches of *sqh_Sqh::GFP kniGal4* control embryos and *sqh_Sqh::GFP kniGal4 Rok::dGBP1* embryos; a-cat::mCherry (red) marks cell junctions. Scale bars, 10 μm or 5 μm on the magnification panel.

For dorsal branch imaging, series of z-stacks was acquired for each embryo at 0.25 μm steps at 3-5 min intervals using Point Scanning Confocal Zeiss LSM880 AiryScan inverted microscope with AiryScan detector (FAST Optimized mode) with the voxel size: 0.0993×0.0993×0.2455 micron^3. Raw images were AiryScan processed with Zen Black software using the “auto” settings to obtain best reconstruction possible.

### Confocal image processing

Z-stack maximum projections were assembled in Fiji (ImageJ)^51^ or OMERO^52^. If needed, signal was enhanced by multiplication i.e., until the Sqh::GFP signal clearly marked the lateral epidermis boundaries. For movies, x-y drift in time was corrected using the StackReg plugin with “rigid body” transformation^53^. Timepoint zero was defined as the beginning of germ-band retraction (trachea imaging) or right before the onset of zippering (70-90 minutes after germband retraction completion) at stage 13/14, for dorsal closure. Figures were prepared using OMERO Figure^52^ and Adobe Illustrator.

### Quantification and exclusion criteria

The number of analyzed embryos of each genotype is shown on the respective figures and indicated in the corresponding text. The following cases were excluded from analysis and were not counted: embryos that were rotated at the time zero or rotated during imaging to an extent that the part of interest was no longer visible/was out of focus; embryos with clear signs of mechanical disruption (pinching of embryo membrane while mounting). For fixed samples, only embryos at stage range indicated below the figure were analyzed. Embryos with clear mechanical disruption signs (independent of the phenotype) or embryos with aberrantly low control mCherry-nls signal were excluded.

### Data availability

The data supporting the findings of this study are available from the corresponding author on request.

## Results

### Design and expression of engineered N-Rok kinase for targeted protein phosphorylation

In order to generate a synthetic kinase that allows for targeted fluorescent fusion protein phosphorylation, we cloned an in-frame fusion of the N-terminal catalytic domain of *Drosophila* Rho-kinase (called N-Rok thereafter) with a small fluorescent protein binder recognizing either GFP or mCherry (Fig. 1c, d, f). We deleted the C-terminal part of Rok, since its removal generates a constitutively active form of the latter^26^. We constructed a series of plasmids to express various combinations of N-Rok and either vhhGFP4 (GFP nanobody)^16^ or 2m22 (mCherry DARPin)^17^ under the control of the UAS-Gal4 system^54^ (Fig. 1d, f and Supplementary Data 1 – FASTA file). In N-RokDead::vhhGFP4 and N-RokDead::2m22, the catalytic domain was rendered inactive by a K116G single amino acid substitution^24^. In a similar way, we engineered a N-Rok::dGBP1 fusion that harbors a destabilized GFP-binding nanobody (dGBP1)^33^, which is stable in the presence of GFP- or its close derivates (Fig. 1e), but degraded in their absence.

In order to generate proof of concept for the engineered kinases, we transfected a stable *Drosophila* S2 cell lines expressing Sqh::GFP (S2 Sqh::GFP)^46^ with pActin_Gal4 and a series of plasmids expressing Rok variants, namely pUASTattB_N-Rok::vhhGFP4-HA, pUASattB_N-RokDead::vhhGFP4-HA, pUASTattB_N-Rok-HA, and pUASattB_N-RokDead-HA. In the interphase, unphosphorylated Sqh localizes mainly to the cytoplasm and its phosphorylation at mitotic entry is known to trigger a cortical recruitment ^55^. To address whether the expression of engineered Rok was sufficient to recruit Sqh::GFP to the cell cortex, where Rok normally exerts its function, we documented the subcellular localization of Sqh::GFP by confocal microscopy in interphasic cells 40 hours after transfection.

In mock transfected cells, the Sqh::GFP signal was predominantly cytoplasmic with no obvious cortical localization (Fig. 2a). In addition to this signal, we observed a prominent cortical enrichment of Sqh::GFP in cells upon the expression of N-Rok::vhhGFP4-HA (Fig. 2a, b). By contrast, cells expressing the non-functional construct (N-RokDead::vhhGFP4-HA) displayed similar Sqh::GFP signals to those seen in mock transfected cells (Fig. 2a, b). Importantly, expression of activated Rok or kinase-dead Rok lacking the fused GFP nanobody (N-Rok-HA or N-RokDead-HA) did not recruit Sqh::GFP at the cortex (Fig. 2a, b).

Of note, expression of N-Rok::vhhGFP4-HA and N-Rok-HA induced cell blebbing and resulted in a high percentage of dead cells, indicating that N-Rok might be toxic in S2 Sqh:: GFP cells, and that only the cells expressing very little active Rok survived. This toxicity, presumably resulting from high expression levels of active Rok with the actin promoter, did not hamper single cell-based analyses (Fig. 2a, b), but prevented us to perform any valuable protein extraction to conduct biochemical analyses of the effects of engineered Rok. In the small fraction of surviving cells expressing N-Rok::vhhGFP4-HA, we detected an enrichment of the Sqh2P signal at the cell cortex using antibody staining, consistent with the higher Sqh::GFP signal at these sites (Fig. 2c). Control cells displayed diffuse Sqh2P signal without enrichment at the cell cortex. A more direct biochemical and mechanical readout of myosin II activation is of importance, as it would more directly validate the relevance of our synthetic kinases approach.

To overcome the limitations of our cell culture system, we generated transgenic *Drosophila* lines using the plasmid constructs mentioned above, such that the expression of synthetic Rok kinase fusion proteins can be controlled with the UAS-Gal4 system^54^. To achieve the same level of expression of the different constructs with the a given Gal4 driver, we used the phiC31 integrase system^45^ and a unique site of integration in an intergenic region on the third chromosome (ZH-86Fb). The first genomic insertions we generated were *N-Rok::vhhGFP4*^ZH-86Fb^ and *N-RokDead::vhhGFP4*^HZ-86Fb^ (Fig. 1d).

### N-Rok::vhhGFP4ZH-86Fb efficiently phosphorylates Sqh::GFP *in vivo*

In order to assess the function of the synthetic Rok kinases in flies, we first investigated whether N-Rok::vhhGFP4^ZH-86Fb^ can regulate Sqh::GFP activity, and whether it can phosphorylate it on serine 21 (Ser19 in vertebrates), the primary activating residue in drosophila (Fig. 1b)^56^. We performed both fluorescence and immunofluorescence analyses on *Drosophila* embryos carrying either *Sqh::GFP (Sqh::GFP* embryos) or *Sqh::mCherry (Sqh::mCherry* embryos), two functional transgenes that can rescue the *sqh^AX3^* null allele in living animals^56–58^. We used *engrailed Gal4* (*enGal4*) to drive the expression of either N-Rok::vhhGFP4^ZH-86Fb^ or N-RokDead::vhhGFP4^ZH-86Fb^ in the posterior compartment of each segment of the embryonic epidermis^59^. For immunostainings, we made use of a rabbit anti phospho-Myosin Light Chain 2 (Ser19) antibody recognizing Drosophila pSqh (#3671, Cell Signaling Technology) and guinea pig anti Sqh1P antibody^47^.

As previously reported^60,61^, actomyosin formed a cable-like structure surrounding the dorsal opening during the stages 13-14 (dorsal closure) in control *Sqh::GFP* embryos (Fig. 2d, white arrow). Expression of N-Rok::vhhGFP4^ZH-86Fb^ in *Sqh::GFP* embryos dramatically altered the distribution of Sqh::GFP, leading to the formation of large foci within the *en* expression domain from late germ band retraction stage onward (stage 12). Sqh::GFP clusters were most prominent in epidermal cells surrounding and contacting the amnioserosa during the dorsal closure stages (Fig. 2d, yellow arrows on the magnified image on the right panel), disrupting the uniform actomyosin cable structure observed in control *Sqh::GFP* embryos (Fig. 2d, white arrow). In contrast, expression of inactive kinase N-RokDead::vhhGFP4^ZH-86Fb^ did not lead to Sqh::GFP clustering (Fig. 2d) nor to a disruption of the uniform actomyosin cable. Of note, we observed Sqh::GFP signal enhancement in the posterior compartments of these embryos (*enGal4* domain); however, this signal was uniformly spread along cell boundaries as normally observed for Sqh::GFP, without the formation of aberrant foci. We assume that this signal enhancement is due to the binding of vhhGFP4 to GFP, as it was previously reported that binding of vhhGFP4 to GFP results in a significant increase in fluorescence signal^22,62^ and we observed similar signal increase upon expression of vhhGFP4 binder alone in *Sqh::GFP* embryos (Supplementary Fig. 1).

In embryos expressing N-Rok::vhhGFP4^ZH-86Fb^, antibody staining revealed that phosphorylated Sqh (p-Sqh) was present in many intense spots overlapping the fluorescent signal emerging from the ectopic Sqh::GFP (n=27/30 embryos included in the analysis), demonstrating efficient Sqh::GFP phosphorylation *in vivo* (Fig. 2d and yellow arrows on the magnified panels on the right). Of note, such pSqh-rich structures were never observed in the adjacent control epidermal stripes. Embryos expressing N-RokDead::vhhGFP4^ZH-86Fb^ (n=22) displayed a diffuse weak p-Sqh signal, as observed in control embryos lacking the synthetic kinase (n=31) (Fig. 2d). Importantly, expression of N-Rok::vhhGFP4^ZH-86Fb^ in the absence of Sqh::GFP (n=13) also showed weak, diffuse p-Sqh signals similar to controls.

To further confirm that the observed potent effects of N-Rok::vhhGFP4^ZH-86Fb^ on Sqh::GFP accumulation and phosphorylation were elicited independently of a protein binder-GFP interaction, we investigated the Sqh::GFP signal and p-Sqh levels in embryos expressing N-Rok::HA^ZH-86Fb^ without the fused nanobody (n=6). In this condition, both the Sqh::GFP and the p-Sqh pattern looked similar to the one seen in control embryos (Fig. 2d, white arrow pointing to the actomyosin cable), indicating that the formation of Sqh::GFP foci and efficient phosphorylation of Sqh is indeed strictly dependent on the binding of the kinase to the fluorescent fusion protein upon the expression of N-Rok::vhhGFP4^ZH-86Fb^. Further evidence for this interpretation was obtained via the expression of N-Rok::vhhGFP4^ZH-86Fb^ in *Sqh::mCherry* embryos, which did neither lead to the formation of Sqh::mCherry foci nor to an alteration of the p-Sqh signal (n=9; add to Fig. 2d).

Taken together, our results show that N-Rok::vhhGFP4^ZH-86Fb^ is able to phosphorylate Sqh::GFP on Ser21, the most important residue for myosin II activation^56^, and indicate that direct interaction between the protein binder and the GFP-target is important for the activity of the kinase domain on the target, thereby allowing for efficient and specific phosphorylation.

### Synthetic Rok kinase modulates mechanical properties of cells through myosin II activity *in vivo*

Phosphorylation of Sqh/Myosin regulatory light chain leads to myosin II activation and the assembly of bipolar myosin filaments that bind F-actin to generate contractility (Fig. 1b) (reviewed in^63^). After confirming that N-Rok::vhhGFP4^ZH-86Fb^ binds to and phosphorylates Sqh::GFP (above), we decided to test whether synthetic Rok was able to specifically modulate mechanical properties of cells through myosin II activity. We focused on dorsal closure, a key morphogenetic process midway through *Drosophila* embryogenesis, where the lateral epidermis from the two sides of the embryo moves up and converges at the dorsal midline to close the epidermal hole, initially covered by amnioserosa cells^60,64^ (Fig. 3a scheme, Fig. 3a “control” and Supplementary Movie 1_Dorsal closure). Dorsal closure represents a well characterized model to study epithelial cell sheet movements and fusion, such as neural tube closure or wound healing in vertebrates, and requires, among other forces, orchestrated actomyosin interplay^65^.

In a similar paradigm as for assessing p-Sqh levels, we used the *enGal4* driver to express N-Rok::vhhGFP4^ZH-86Fb^ in a striped pattern in the embryonic epidermis and analyzed its effects on epithelial cell behavior. We performed live imaging in either *Sqh::GFP* or *Sqh::mCherry* embryos, following their development from dorsal closure (stage 13-15) onward.

In contrast to control *Sqh::GFP* embryos, in which dorsal closure occurred normally (n=12/14) (Fig. 3a and Supplementary Movie_1_Dorsal closure), expression of N-Rok::vhhGFP4^ZH-86Fb^ in the *Sqh::GFP* background led to abnormal dorsal closure (n=13) (in most severe cases resulting in a dorsal open phenotype) due to the formation of Sqh::GFP foci and additional cable structures under tension in the expression domain of N-Rok::vhhGFP4^ZH-86Fb^ (Fig. 3a and Supplementary Movie 1_Dorsal closure). These structures resulted in a disruption of the actomyosin cable and in local invaginations (presumably due to ectopic pulling forces within the entire tissue) and deformation of the entire epidermis, which prevented normal closure. Contrary to N-Rok::vhhGFP4^ZH-86Fb^, *Sqh::GFP* embryos in which N-RokDead::vhhGFP4^ZH-86Fb^ was expressed (n=8) displayed a well-discernible actomyosin cable around the amioserosa, and dorsal closure proceeded normally (Fig. 3a and Supplementary Movie 1_Dorsal closure). As described above for the fixed samples, we observed a uniform Sqh::GFP signal enhancement on cell boundaries in the expression domain of N-RokDead::vhhGFP4^ZH-86Fb^, an effect most likely due to binding of vhhGFP4 to GFP^22,62^. In the embryos expressing N-Rok::HA^ZH-86Fb^ without the nanobody (n=8/9), the actomyosin cable formed as in wildtype embryos, accompanied by normal dorsal closure (Supplementary Fig. 2 and Supplementary Movie 2_Dorsal closure); no uniform Sqh::GFP signal enhancement was seen in these conditions. Most of the embryos expressing N-Rok::vhhGFP4^ZH-86Fb^ without GFP::Sqh closed normally (n=11/14) (Fig. 3a and Supplementary Movie 1_Dorsal closure). As expected, expression of N-Rok::vhhGFP4^ZH-86Fb^ in *Sqh::mCherry* embryos (n=4) also lead to normal closure (Supplementary Fig. 2 and Supplementary Movie 3_Dorsal closure).

The phenotypes described in this and in the following section were fully penetrant and occurred in the presence and in the absence of a wild type *sqh* gene, rendering the sexing of the embryos unnecessary. Interestingly, we noticed during the course of our experiments that the phenotype was more severe when embryos were mounted by gluing them to the coverslip (dorsal open, see Fig. 3b), while a slightly milder phenotype was seen when the embryos were imaged without gluing, in a glass-bottom dish (aberrant closure, see Fig. 3a) (see Materials & Methods section for procedure details). We presume that these differences result from differences of external tension induced by each of the mounting techniques. Only for Fig. 3b, c and corresponding Supplementary Movie 4_Dorsal closure, the embryos were imaged with gluing to the coverslip technique; all the other presented embryos were imaged on a glassbottom dish.

We conclude from these results that N-Rok::vhhGFP4^ZH-86Fb^ efficiently targets and phosphorylates Sqh::GFP but not Sqh::mCherry nor endogenous Sqh, and affects mechanical properties of cells through myosin II activity.

### Comparison of engineered Rok with previously available tools for direct activation of myosin II

In order to compare N-Rok::vhhGFP4^ZH-86Fb^ to the already published effectors that are known to activate myosin II, we used the *enGal4* driver and embryonic dorsal closure to benchmark N-Rok::vhhGFP4^ZH-86Fb^ and a few chosen activators. Two constitutively active forms of Rok (Rok-CAT and Rok-cat) have been generated, and several independent random genomic insertions of *Rok-CAT*^24^ and *Rok-cat*^27^ generated by P element mediated transformation are available. In this work, we only report results obtained with *Rok-CAT^3.1^* and *Rok-cat^T2A^*, the insertions that yielded the highest levels of expression of these transgenes^24,27^. In addition, we compared N-Rok::vhhGFP4^ZH-86Fb^ with Sqh^E20E21^, a phosphomimetic that emulates diphosphorylated Sqh^24^, as well as with ctMLCK, a constitutively active form of chicken Myosin light-chain kinase, which is known to phosphorylate Sqh^66^. Table 1 summarizes the different effectors we tested, the genetic backgrounds in which we performed those tests, and their effect on dorsal closure.

**Table 1:** Benchmarking of the direct myosin II effectors available in *Drosophila*. Dorsal closure in embryos and global viability were used as a readout of myosin II activation in the *en* expression domain. Both the effectors and the full genotypes of the tested embryos are indicated to highlight all the components that the methods require. For every condition, more than 20 embryos were assessed. See text for details.

| effector | genotype | dorsal closure phenotype | viability |
| --- | --- | --- | --- |
| Rok-cat<sup>T2A</sup> | Sqh::GFP, enGal4 / + ; Rok-cat<sup>T2A</sup> / + | aberrant | larval lethal |
| Rok-CAT<sup>3.1</sup> | Sqh::GFP, enGal4 / + ; Rok-CAT<sup>3.1</sup> / + | wild type | viable |
| N-Rok::vhhGFP4<sup>ZH-86Fb</sup> | Sqh::GFP, enGal4 / + ; N-Rok::vhhGFP4<sup>ZH-86Fb</sup> / + | aberrant/open | embryonic lethal |
| N-Rok::HA<sup>ZH-86Fb</sup> | Sqh::GFP, enGal4 / + ; N-Rok::HA<sup>ZH-86Fb</sup> / + | wild type | reduced viability |
| N-Rok::vhhGFP4<sup>Vi</sup> | Sqh::GFP, enGal4 / + ; N-Rok::vhhGFP4<sup>Vi</sup> / + | aberrant | embryonic lethal |
| Sqh<sup>E20E21</sup> | enGal4 / Sqh<sup>E20E21</sup> | wild type | viable |
| ctMLCK | Sqh::GFP, enGal4 / + ; ctMLCK / + | aberrant or normal but slower | larval lethal |
| NSlmb::vhhGFP4 | flw::GFP, enGal4 / + ; NSlmb::vhhGFP4 / + | wild type | viable |

*Sqh::GFP* embryos expressing Rok-CAT (n=5) did not show any dorsal open phenotype and were viable (Fig. 3c and Supplementary Movie 4_Dorsal closure). In contrast, expression of Rok-cat (n=4) in *Sqh::GFP* embryos was larval lethal and resulted in either a mild stripe misalignment or a more severely aberrant dorsal closure with ectopic clusters of tension (Fig. 3c and Supplementary Movie 4_Dorsal closure). Expression of Sqh^E20E21^ under the control of *enGal4* (n=5, Supplementary Fig. 2, Supplementary Movie 2_Dorsal closure) or under the endogenous promoter of *sqh*^24^ were both viable conditions that produced adult flies (Table 1). *Sqh::GFP* embryos expressing ctMLCK showed varying effects on dorsal closure: the process was either aberrant (n=5/8) or successfully completed but at a lower speed than observed normally (n=3/8) (Fig. 3c and Supplementary Movie 4_Dorsal closure).

During the course of our work, studies from the Sanson lab reported hyperactivation of Myosin II via knockdown of the Myosin II phosphatase Flapwing (Flw)^67^. They used NSlmb::vhhGFP4 (deGradFP^68^) to achieve efficient protein knockdown of Flw::GFP. However, expression of NSlmb::vhhGFP4 with the *enGal4* driver in *flw*::GFP embryos (this work) produced adult flies, indicating complete dorsal closure (Table 1) and implying less potent Myosin II activation then achieved with the use of N-Rok::vhhGFP4^ZH-86Fb^.

Overall, N-Rok::vhhGFP4^ZH-86Fb^ turned out to be the most efficient specific myosin II activator among the tools we tested and the assays we used, together with ctMLCK^66^, which, at least in some embryos, resulted in similarly strong dorsal closure phenotype.

### Evaluation of the general toxicity of N-Rok::vhhGFP4^ZH-86Fb^ in *Drosophila*

Biochemical and cell culture studies reported several potential substrates for Rok/ROCK (reviewed in^69^). We presumed that binder-dependent interaction of N-Rok::vhhGFP4^ZH-86Fb^ with Sqh::GFP generates a more specific tool for directed myosin II targeting than the previously available, constitutively active effectors (Rok-CAT and Rok-cat)^24,27^. However, the specificity of N-Rok::vhhGFP4^ZH-86Fb^ toward a GFP fusion target is unlikely to be absolute, in particular due to the high expression when using the Gal4 system, and it is important to discriminate between myosin II-dependent effects and the effects of other possible targets on the resulting phenotype.

To evaluate the general toxicity of N-Rok::vhhGFP4^ZH-86Fb^ in *Drosophila*, and to compare it to the toxicities of Rok-CAT and Rok-cat, we used different Gal4 drivers and expressed them in flies that neither carried a *Sqh::GFP* nor any other GFP fusion target transgene. Doing so, we analyzed the viability and, for viable conditions, the adult fly phenotypes generated by the expression of either N-Rok::vhhGFP4^ZH-86Fb^, N-RokDead::vhhGFP4^ZH-86Fb^, Rok-CAT or Rok-cat in different tissues and with different efficiency (Supplementary Table 1).

Overall, the phenotypes increased in severity according to the potency of the Gal4 drivers used. Importantly, *N-Rok::vhhGFP4*^ZH-86Fb^ turned out to induce severe phenotypes in combination with all Gal4 drivers tested (Supplementary Table 1). These phenotypes were similar to the phenotypes produced by *Rok-cat^T2A^* in combination with the same Gal4 drivers (^27^ and Supplementary Table 1). As shown above (Fig. 3a), embryos expressing *N-Rok::vhhGFP4*^ZH-86Fb^ in the *en* stripes without a GFP-target closed mostly normally (n=11/14) and did not show increased p-Sqh levels (Fig. 2d); nevertheless, they were embryonic/larvae lethal. Of note, *N-RokDead::vhhGFP4*^ZH-86Fb^ expressed without a GFP-target did not cause any detrimental phenotype (Supplementary Table 1), suggesting that the GFP-independent effect of *N-Rok::vhhGFP4*^ZH-86Fb^ was due to its active kinase domain.

We noticed that *Rok-CAT^3.1^* induced milder phenotypes than *Rok-cat^T2A^*, despite little differences between their sequences^24,27^. This was also true with regard to dorsal closure, which was mostly aberrant for *Rok-cat^T2A^* and normal for *Rok-CAT^3.1^* (described above, and Fig. 3c). Hence, we assumed that the differences of general toxicity between the constitutively active *N-Rok::vhhGFP4*^ZH-86Fb^, *Rok-cat*^T2A^, and *Rok-CAT*^3.1^ are most likely due to the different expression levels from the different genomic insertion sites.

### Generation of a less toxic N-Rok::vhhGFP4^Vi^ genomic insertion

In order to validate whether the observed differences in the overall toxicity of the tested genomic insertions are due to divergent expression levels of constitutively active Rok, we tried to generate insertions of *N-Rok::vhhGFP4* that would display as little toxicity as possible when combined with Gal4 drivers in the absence of a GFP-fusion target. We generated 25 independent random genomic insertions of *N-Rok::vhhGFP4* by P-element mediated transformation^48^ and screened for an insertion that would be viable and fertile in combination with the *enGal4* driver.

Only one insertion (called *N-Rok::vhhGFP4*^Vi^, for “viable”) among the 25 lines we generated was viable and fertile in combination with *enGal4*. Since expression of *enGal4* is turned on in the posterior compartment of each segment of the epidermal structures, including the wings, we analyzed the wings of *N-Rok::vhhGFP4*^Vi^ flies. We solely noticed a wing phenotype consisting of partially missing crossvein and a slight but significant reduction of the posterior compartment relative to the total wing area (Supplementary Fig. 3). Interestingly, the partially missing crossvein phenotype is reminiscent of the phenotype due to ectopic activation of Rho1 during wing development, which is in line with N-Rok::vhhGFP4^Vi^ mildly activating myosin II^27,70^. *N-Rok::vhhGFP4*^Vi^ did not show any phenotype when combined with *elavGal4*, with *69BGal4* or with *daGal4* driver; in contrast, it turned out to be lethal in combination with *tubGal4*, which has a strong ubiquitous expression pattern (Supplementary Table 1). Taken together, these results suggest that the *N-Rok::vhhGFP4*^Vi^ line allows to achieve expression levels of N-Rok::vhhGFP4 low enough to prevent apparent off-target effects in *Drosophila* with most Gal4 drivers.

To test whether *N-Rok::vhhGFP4*^Vi^ can be used to express enough N-Rok:: vhhGFP4 to act as a useful source of an engineered kinase, we again used the *enGal4* driver and dorsal closure as an assay system to assess the effects on myosin II contractility. Similarly to N-Rok::vhhGFP4^ZH-86Fb^, expression of N-Rok::vhhGFP4^Vi^ in *Sqh::GFP* embryos resulted in the formation of large Sqh::GFP foci in every other segment of the embryonic epidermis that were most prominent around the dorsal hole (Fig. 4b). As observed for N-Rok::vhhGFP4^ZH-86Fb^ (Fig. 2d), those large Sqh::GFP foci overlapped with p-Sqh clusters (n=14/15, Fig. 4b). In the same paradigm as for *N-Rok::vhhGFP4*^ZH-86Fb^, we performed live imaging in *Sqh::GFP* embryos expressing N-Rok::vhhGFP4^Vi^ with *enGal4*. This condition led to abnormal dorsal closure (n=10) and was lethal, similarly to *N-Rok::vhhGFP4*^ZH-86Fb^ (Fig. 4c and Supplementary Movie 5_Dorsal closure).

We conclude from these results that *N-Rok::vhhGFP4*^Vi^ allows to express a synthetic kinase whose concentration is both high enough to phosphorylate a GFP-fusion version of Sqh, and low enough to prevent deleterious effects in flies deprived of a GFP target.

### Generation of a destabilized GFP-binder kinase

During the course of our studies, the group of Cepko reported the isolation and characterization of a destabilized GFP-binding nanobody (dGBP1)^33^. dGBP1 fusion proteins are only stable in the presence of a GFP (or its close derivates), both *in vitro* and *in vivo;* in the absence of a GFP fusion protein, dGBP1 is destabilized and the entire dGBP1-fusion protein is degraded by the ubiquitin proteasome system, with no signs of aggregation in cells^33^ (Fig. 4a).

In order to further minimize the potential off-target activity of our synthetic N-Rok kinase in the absence of a GFP-target, we generated flies carrying *N-Rok::dGBP1* (Fig. 1e) in the same intergenic region on the third chromosome (ZH-86Fb) as used to generate *N-Rok::vhhGFP4*^ZH-86Fb^ flies. When we expressed N-Rok::dGBP1^ZH-86Fb^ within the *en* domain in *Sqh::GFP* embryos, we observed spots of high p-Sqh levels overlapping with large Sqh::GFP foci expressing in every other segment (Fig. 4d). Despite large Sqh::GFP clusters, p-Sqh foci were less prominent than those observed for N-Rok::vhhGFP4^Vi^ (Fig. 4a) or N-Rok::vhhGFP4^ZH-86Fb^ (Fig. 2d); however, they were still clearly distinguishable in most embryos (n=29/35). In contrast, the embryos expressing N-Rok::dGBP1^ZH-86Fb^ alone displayed a dim, uniform p-Sqh signal (n=4, Fig. 4d).

Along the same lines as for N-Rok::vhhGFP4^Vi^ (Fig. 4b) and N-Rok::vhhGFP4^ZH-86Fb^ (Fig. 3a), we performed live imaging of dorsal closure in *Drosophila* embryos carrying *Sqh::GFP* and *N-Rok::dGBP1*^ZH-86Fb^ expressed in the *en* domain (n=10). As expected, we observed the formation of ectopic cable structures that disrupted the uniform actomyosin cable string (Fig. 4e and Supplementary Movie 5_Dorsal closure), leading to the abnormal closure and embryonic/early larvae lethality. Most importantly, embryos expressing only N-Rok::dGBP1^ZH-86Fb^ in the *en* domain without the GFP target exhibited normal closure, did not show any obvious developmental defects and proceeded to adulthood (n=12, Fig. 4e and Supplementary Movie 5_Dorsal closure).

Our data shows that N-Rok::dGBP1^ZH-86Fb^ phosphorylates Sqh::GFP and efficiently modulates mechanical properties of cells through myosin II, similarly to N-Rok::vhhGFP4^Vi^ and N-Rok::vhhGFP4^ZH-86Fb^. More importantly, expression of N-Rok::dGBP1^ZH-86Fb^ did not show any obvious deleterious effects in flies devoid of a GFP target (Fig. 4e). We presume that this is due to the degradation of the N-Rok::dGBP1 fusion protein in the absence of a dGBP1 cognate antigen, thereby restricting the kinase activity to the time and place where a GFP-target is present.

### Nanobody-based engineered kinases are modular tools

Nanobody-based engineered kinases would display an extra degree of modularity if one could target the kinase domain to other fluorescent proteins than GFP and its close derivatives. To that aim, we first generated two novel genomic insertions in ZH-86Fb: *N-Rok::2m22*^ZH-86Fb^ and *N-RokDead::2m22*^ZH-86Fb^. *N-Rok::2m22*^ZH-86Fb^ and *N-RokDead::2m22*^ZH-86Fb^ are similar to *N-Rok::vhhGFP4*^ZH-86Fb^ and *N-RokDead::vhhGFP4*^ZH-86Fb^, respectively, but a DARPin specifically recognizing mCherry^17^ replaces the nanobody binding to GFP (Fig. 1f).

Similarly to our assays for N-Rok::vhhGFP4^ZH-86Fb^ in *Sqh::GFP* embryos, we tested whether N-Rok::2m22^ZH-86Fb^ can specifically modulate myosin II activity through the phosphorylation of Sqh::mCherry. As expected, expression of N-Rok::2m22^ZH-86Fb^ in *Sqh::mCherry* embryos with the *enGal4* driver led to the formation of large Sqh::mCherry foci in every other segment of the embryonic epidermis, with a prominent p-Sqh signal corresponding to the foci (n=8, Supplementary Fig. 4). In contrast, *Sqh::GFP* embryos did not show any clusters in the *en* domain upon N-Rok::2m22^ZH-86Fb^ expression and exhibited diffuse and wild type-like p-Sqh staining (n=8, Supplementary Fig. 4).

We next performed live imaging experiments in *Sqh::mCherry* embryos. Again, we used *enGal4* to drive the expression of *N-Rok::2m22*^ZH-86Fb^ and analyzed dorsal closure. In contrast to the control *Sqh::mCherry* embryos, which closed normally (n=8, Supplementary Fig. 2 and Supplementary Movie 3_Dorsal closure), expression of N-Rok::2m22^ZH-86Fb^ in the presence of Sqh::mCherry led to abnormal closure (n=4) as well as to the formation of cable structures under tension (Supplementary Fig. 2 and Supplementary Movie 3_Dorsal closure) and was larval lethal. Importantly, in *Sqh::mCherry* embryos expressing *N-RokDead::vhhGFP4*^ZH-86Fb^ dorsal closure proceeded normally (n=5, Supplementary Fig. 2 and Supplementary Movie 3_Dorsal closure).

We conclude from these results that N-Rok::2m22^ZH-86Fb^ can efficiently and specifically phosphorylate Sqh::mCherry modulating myosin II activity in a way similar to N-Rok::vhhGFP4 in *Sqh::GFP* embryos.

### Can engineered Rok kinase target GFP-proteins that are not its natural target?

We have shown that a synthetic Rok kinase activates myosin II through the phosphorylation of Sqh::GFP, thereby influencing tissue mechanics *in vivo*. We next sought to test whether N-Rok::vhhGFP4 can phosphorylate and/or activate other GFP-fusion proteins, which are not a direct target of Rok in cells.

We first used a *Tkv::YFP* line (a gift from Georgios Pyrowolakis) and expressed N-Rok::vhhGFP4^Vi^ in the *en* compartment of the embryos. Thickveins (Tkv) is a receptor of the Decapentaplegic (Dpp) morphogen^71^. Upon binding of Dpp, Tkv is activated by phosphorylation of specific cytoplasmic S/T residues by its co-receptor Punt, eventually resulting in Mad phosphorylation^72–76^. We therefore reasoned that Tkv could potentially be a substrate for our engineered N-Rok::vhhGFP4 (S/T kinase) and used p-Mad activation levels as a readout. In these conditions, striped expression of N-Rok::vhhGFP4^Vi^ in *Tkv::YFP* embryos had no effect on pMad staining levels (Supplementary Fig. 5a) nor on embryo dorsal closure (data not shown). Moreover, using another Tkv line, we verified that expression of Tkv::GFP and N-Rok::vhhGFP4 in the engrailed domain did not influence p-Sqh levels (n=5) (Supplementary Fig. 5b).

Rok/ROCK has been shown to interact with the cadherin complex through direct binding to p120 catenin and to influence adherens junctions stability^77^. E-cadherin is phosphorylated on a few serine residues and this phosphorylation is required for binding to β-catenin and the stability of this complex^78,79^. We therefore decided to use *DE-Cadherin::GFP^80^* embryos to ask if our synthetic Rok kinase could act on DE-Cadherin::GFP and cause any obvious developmental effects. We used the *enGal4* driver to express N-Rok::vhhGFP4^Vi^ and focused on dorsal closure and viability. We did not observe any detrimental effect (n=5/6 embryos) in the presence of the activated kinase and this condition was viable, similarly to embryos harboring the balancer instead of kinase-containing chromosome (n=5/6 embryos) (Supplementary Fig. 5c and Supplementary Movie 6_Dorsal closure) or *yw* control flies crossed with Cadherin::GFP (8/8 embryos) (data not shown). Of note, we observed enhanced GFP signal in the *en* domain, inferring N-Rok::vhhGFP4^Vi^ binding to GFP, according to the previously reported increase in fluorescence signal upon binding of vhhGFP4 to GFP (see Fig. 2d, Supplementary Movie 6_Dorsal closure and ^22,62^).

We assumed that since Rok kinase normally exerts its function in the vicinity of cell membrane^81^, expression of N-Rok::vhhGFP4^Vi^ in the presence of GFP-tagged DE-Cadh may increase the chance of unspecific phosphorylation of the untagged Sqh protein present in the cell cortex. The increased GFP signal in the *en* domain observed in the above experiment implies that N-Rok::vhhGFP4^Vi^ binds DE-Cadh::GFP, suggesting it is enriched in close proximity of cell membrane. We therefore sought to examine if endogenous Sqh becomes a target when the engineered Rok kinase is brought in close proximity through neighboring GFP-binder interactions. We examined the levels of p-Sqh in *DE-Cadh::GFP* embryos expressing N-Rok::vhhGFP4^Vi^ with the *enGal4* driver. We did not observe a p-Sqh signal increase in this setup (as observed for N-Rok::vhhGFP4^Vi^ in the presence of Sqh::GFP) (n=6, Supplementary Fig. 5d).

These results suggest that GFP-fusion kinases have to be brought into direct contact to their target proteins in order to affect their phosphorylation; simply being enriched in the same cellular region does not appear to be sufficient to direct their activity efficiently towards a given target protein.

### The tracheal system of Drosophila embryo develops abnormally upon excessive myosin II phosphorylation by engineered Rok kinase

We next wanted to use the engineered Rok kinase to study the effects of actomyosin activation on cell migration and cell rearrangements during branch formation in the tracheal system, the respiratory organ of a fly. In the *Drosophila* embryo, the tracheal system develops from 10 bilaterally symmetrical clusters of ectodermal cells. Cells in each cluster invaginate and migrate to form one tracheal segment that further connects with neighboring segment to form the stereotypical network of tubes ^82–85^. Remarkably, cells do not divide during this process, but arrange into the complex branched structure via cell migration, cell shape changes and cell-cell neighbor exchange. During the formation of the dorsal branch (DB), individual DBs initially elongate as a cluster of cells, led by migrating distal-most cells at the front (tip cells), which induces tension on stalk cells trailing in the back, which ultimately results in stalk cell intercalation (SCI) and elongation of stalk cells ^82,86^. SCI occurs in all primary branches except for the ones that will eventually form the dorsal trunk, the largest tube in the tracheal system ^86–88^. Surprisingly, and in contrast to many other morphogenetic events of tissue elongation through cell intercalation ^34–43^, DB cell intercalation occurs in the absence of MyoII function; intercalation is thus a passive event with regard to force generation in stalk cells, and it is possible that stalk cells need to downregulate actomyosin activity to give in to the tension created by the migration of tip cells ^89^.

To test whether the activation of MyoII in tracheal cells would lead to a failure in the release of tension and consequently to the inhibition of proper trachea development, we expressed the various variants of synthetic N-Rok using the pan-tracheal *breathless-Gal4* (*btlGal4*) driver in the *sqh_Sqh::GFP* background. We first tested whether engineered Roks phosphorylate Sqh::GFP specifically in the developing trachea. Expression of all three active N-Rok variants (N-Rok::vhhGFP4^ZH-86Fb^, N-Rok::vhhGFP4^Vi^ and N-Rok::dGBP1^ZH-86Fb^) resulted in the formation of large Sqh::GFP foci in tracheal cells that represented highly phosphorylated Sqh as confirmed by pShq antobody staining (n=7/8, n=7, n=4/6, respectively, Fig. 5a). In most cases, we observed a clustering of groups of tracheal cells and a discontinued tracheal network, as shown for N-Rok::vhhGFP4^ZH-86Fb^ and N-Rok::dGBP1^ZH-86Fb^, and visualized by mCherry-nls marking tracheal cell nuclei (yellow arrows in Fig. 5a). We presume that these clusters arise from the accumulation of tracheal cells around the Sqh::GFP foci as a result of increased tension, “holding” the clusters of cells together and preventing them from proper migration and intercalation. In the most severe cases, almost no cells escaped from the “rosettes” around the Sqh::GFP foci and the whole cluster of cells were migrating together, without showing any signs of branch formation (see N-Rok::vhhGFP4^Vi^ in Fig. 5a). We noted a variety of phenotypes ranging from very severe (cells forming compacted rosettes around the Sqh::GFP clusters) to slightly milder (several cells escaping from the rosettes and forming branches). Nevertheless, in all cases we observed embryonic/early larval lethality. The images on Fig. 5a were chosen to represent the strongest observed phenotype captured at a given stage (14-16) and in an adequate positioning of the embryo in the specimen. In contrast, embryos expressing inactive N-RokDead::vhhGFP4^ZH-86Fb^ variant did not show clusters of Sqh::GFP nor p-Sqh foci (n=6); tracheal cells migrated out and formed stereotypic branches, similar to what is seen in wild type embryos (Fig. 5a), and this condition was viable.

To better understand the timing and cell behaviors in the context of constitutive activation of myosin II by N-Rok in tracheal cells, we next performed live imaging in *Sqh::GFP* embryos expressing engineered Rok variants in the *btl* expression domain. Importantly, for all active N-Rok variants, we noticed Sqh::GFP clusters at the invaginating tracheal placodes already around stage 11 (yellow arrows in Fig. 5b and Supplementary Movie 7_Trachea development). Sqh::GFP clusters were most prominent for N-Rok::vhhGFP4^ZH-86Fb^ and for N-Rok::vhhGFP4^Vi^ (n=9 for both), and somewhat smaller for N-Rok::dGBP1^ZH-86Fb^ (n=16). As a result, tracheal cells were unable to migrate away from the placode but rather formed rosettes around the Sqh::GFP foci, leading to a dramatically disorganized and fragmented tracheal architecture (Fig. 5b, asterisks and Supplementary Movie 7_Trachea development). Again, as described above for the fixed samples, we noticed a range of phenotypes that differed in how many cells escaped the Sqh::GFP clusters and migrated out forming aberrant, yet distinct dorsal and ventral branches, or in whether entire rosettes migrated together. In contrast, most *Sqh::GFP* embryos expressing the inactive kinase N-RokDead::vhhGFP4^ZH-86Fb^, as well as simultaneously imaged control embryos, developed a morphologically normal tracheal network (n=8/9 and n=7/9, respectively, Fig. 5b Supplementary Movie 7_Trachea development) and this condition was viable.

We conclude from these experiments that tracheal-specific phosphorylation of Sqh::GFP by synthetic N-Rok variants strongly perturbs cell migration from early stages of trachea development onwards, resulting in dramatic phenotypes; disrupted primary branch formation and aberrant trachea development leading to embryonic/early larvae lethality. These results suggest that finely tuned levels of active myosin II are crucial for proper trachea morphogenesis.

### Effect of actomyosin activation on cell intercalation and dorsal branch elongation

Since the use of a pan-tracheal driver (*btl*Gal4) to express active N-Rok kinase resulted in robust Sqh::GFP phosphorylation and cell clustering already at early stages of tracheal development (see above), this scenario prevented us from studying cell intercalation and adherens junction remodeling in developing branches. To take a closer look at these processes in the context of excessive actomyosin activity, we switched to a *knirps* (*kniGal4*) driver line, which activates expression in dorsal and ventral branches, but not in the dorsal trunk ^90^.

At the onset of branch elongation (stage 13/14), stalk cells in dorsal branches are in a side-by-side arrangement; as the branch elongation progresses and cells rearrange, intercellular AJs of adjacent cells are progressively converted into autocellular AJs as the stalk cells reach around the lumen and seal the lumen tube with these newly formed AJs (Fig. 6a) ^86,91^. By stage 16, when SCI is completed, and one of the tip cells meets with the contralateral branch (fusion cell) and the other one becomes a terminal cell with long cellular extensions, stalk cells are in an end-to-end arrangement and show characteristic pattern of lines and small rings of AJs, corresponding to auto- and intercellular AJs, respectively ^92^.

To test the hypothesis that actomyosin activity might need to be low in stalk cells in order not to produce forces counteracting tip cell migration and DB cell intercalation, we used our engineered Rok kinase to look at branch elongation and the stereotypic junction formation in developing dorsal branches. Since the data shown above demonstrates that N-Rok::dGBP1^ZH-86Fb^ phosphorylates Sqh::GFP and efficiently modulates mechanical properties of cells through myosin II (yet at slightly lower levels than N-Rok::vhhGFP4^ZH-86Fb^ or N-Rok::vhhGFP4^Vi^), and, most importantly, that its expression is innocuous in flies devoid of a GFP target, we used this variant synthetic kinase for the following experiments.

We first examined the influence of actomyosin activation in dorsal branches on filopodia formation and cell migration in time-lapse imaging using a LifeAct-mRuby reporter to visualize pools of F-actin. Expression of the engineered Rok kinase N-Rok::dGBP1^ZH-86Fb^ in the *kni* expression domain in *Sqh::GFP* embryos resulted in the formation of Sqh::GFP clusters from stage 13/14 onwards in the areas around cell junctions in the stalk cells and in between the two tip cells (Fig. 6b). Such Sqh::GFP foci were never observed in the control embryos in the absence of N-Rok::dGBP1^ZH-86Fb^ expression. Similarly to the control embryos, Sqh::GFP embryos expressing engineered N-Rok kinase in the *kni* domain formed filopodia and showed high filopodia activity in the tip cells. In both classes of embryos, in most branches stalks cells trailed after the filopodia-rich tip cells, leading to the elongation of the dorsal branches (n_branches_= 102/104 for dGBP1, from n_embryos_=35 and n_branches_= 46/46 for control, from n_embryos_=15, on average 3 central branches per embryo were analyzed; Fig. 6b and Supplementary Movie 8_Branch elongation). Therefore, expression of engineered Rok kinase N-Rok::dGBP1^ZH-86Fb^ with *kniGal4* driver did neither preclude filopodia formation nor cell migration of tip cells.

However, many branches expressing N-Rok::dGBP1^ZH-86Fb^ displayed several defects, most commonly: 1) Some tip cells showed aberrant behavior in keeping persistent contact with adjacent branches (n_branches_=16/104, arrow, Fig. 6b and Supplementary Movie 8_Branch elongation); 2) In some cases, tip cells detached from stalks cells (n_branches_=10/104; arrowheads, Fig. 6b and Supplementary Movie 8_Branch elongation); 3) In a few cases, after initial branch elongation, we observed clustering of tip and stalk cells accompanied by the branch shortening (n_branches_=5/104). As a result, some branches did not establish proper 1:1 contact with the opposite partner branch from the other side of the embryo. Instead, DBs connected only with a neighbor branch or clustered with the adjacent branch and contacted together the contralateral partner branch or – in the case of branch shortening, no contact with the contralateral partner was established. In the control embryos, most of the branches analyzed migrated as expected (n_branches_=44/46) and, after reaching the midline, contacted their contra-lateral partners in a 1:1 ratio (Fig. 6b, last timepoint and Supplementary Movie 8_Branch elongation).

We next performed anti-DE-cadherin staining to investigate cell intercalation and autocellular junction formation in stage 16 embryos. Before intercalation, dorsal branches expressing engineered Rok kinase displayed prominent Sqh::GFP foci in the cluster of cells giving rise to the emerging branch (Fig. 7a, stage 14, arrows). As the branches elongated, the foci moved together with the intercalating stalk cells, eventually persisting predominantly at the sites of the intercellular junctions (Fig. 7a, stage 15-16, arrows). At stage 16, *Sqh::GFP* embryos expressing engineered N-Rok kinase showed characteristic pattern of AJs similarly to the control embryos, indicating correct cell intercalation despite the presence of ectopically activated actomyosin foci (n_branches_=23/26 from n_embryos_=13 for N-Rok::dGBP1^ZH-86Fb^ and n_branches_=14/14 from n_embryos_=6 for the control; on average 2 branches per embryo were analyzed, Fig. 7a, stage 16). Co-staining of the embryos with Vermiform, a marker for the tracheal lumen, revealed that in the vast majority of the imaged embryos (add), the lumen was properly formed both in the dorsal trunk (where no synthetic kinase was expressed), as well as in the kinase-expressing branches (data not shown).

In order to follow cell intercalation in the context of excessive actomyosin activation in more detail, we performed live-imaging in *Sqh::GFP* embryos expressing fluorescent α-Catenin (a-cat::mCherry) to visualize cell junctions (Supplementary Movie 9_Intercalation_control). We used again the *kni* driver to express N-Rok::dGBP1^ZH-86Fb^. In the course of the experiments with *UAS_a-cat::mCherry* stock, we noticed background expression of a-cat::mCherry in the epidermis, regardless of the driver used, and even in the absence of a driver. However, we verified the presence of UAS sequence with a robust signal increase in posterior domain of each segment with the use of *en* driver (data not show). Therefore, the observed background of a-cat::mCherry expression, although limiting a clearer view on the imaged embryos (especially at the beginning of branch emerging), did not prevent the use of this line for studying intercalation in dorsal branches via live-imaging.

In line with the live-imaging experiments using LifeAct-mRuby (see above), we observed Sqh::GFP clusters from stage 13/14 onwards upon expression of engineered kinase N-Rok::dGBP1^ZH-86Fb^ in the *kni* domain. These clusters moved along with the cells in elongating branches, finally accumulating mostly at intercellular AJs, including prominent foci at the tip of the branch (in between the two tip cells) (Fig. 7b, arrows and Supplementary Movie 10_Intercalation_Rok_dGBP1). In most cases, however, we observed adequate AJs rearrangements, reflecting two migrating tip cells and stalk cell intercalation and elongation (n_branches intercalated_=35/36 from n_embryos_=14 for N-Rok::dGBP1^ZH-86Fb^ and n_branches_=32/32 from n_embryos_=12 for the control; on average 2-3 branches per embryo were analyzed, Fig. 7b, Supplementary Movie 9_Intercalation_control and Supplementary Movie 10_Intercalation_Rok_dGBP1).

We therefore conclude that excessive phosphorylation of Sqh::GFP by synthetic N-Rok::dGBP1^ZH-86Fb^ specifically in the *kni* domain during dorsal branch formation did not preclude filopodia activity in tip cells, allowing for cell migration, branch elongation and stalk cells intercalation in most branches, despite causing aberrant behavior/contacts establishment between some of the tip cells. In most cases, autocellular junction were properly formed, although in some cases the regular spatial arrangement of stalk cells was somewhat disrupted or – in most severe cases – parts of the cells detached from the rest of the branch disabling resulting in aberrant branch network.

## Discussion

In this study, we generated and characterized a series of genetically encoded synthetic Rok kinases using protein binders, and provide evidence that such constructs can be used as valuable tools that allows for efficient phosphorylation of a specific target *in vivo* in a tissuespecific manner. We show that tissue-specific expression of an engineered Rok-protein binder leads to efficient phosphorylation of the fluorescently tagged substrate protein Sqh and increases actomyosin contractility, resulting in aberrant cell behavior. While we have studied these engineered Rok kinases in two morphogenetic processes in the developing fly embryo, we assume that this tool can be adapted to other kinases and targets in various eukaryotic genetic systems; however, this will require careful testing for each kinase used.

Protein binder-based approaches for directing a given enzymatic function to a specific, single target open new avenues for unravelling protein-protein interactions in a given cell type and for manipulating them in a desired way, both for developmental studies, for designing synthetic biology systems or eventually for therapeutic goals. Similar nanobody-based paradigm has been recently applied to induce glycosylation and deglycosylation of desired target proteins in cells^93,94^. Furthermore, in the elegant work by Karginov et al., a rapamycin-regulated engineered Src kinase was used in cultured cell to dissect the effect of Src-dependent phosphorylation of its two substrates, FAK and p130Cas (containing a modified rapamycin– binding domain insertion), on cell morphology and filopodia formation^95^.

Activation of myosin II-dependent contractility has been studied in various biological systems in the context of shaping cells and tissues during development, for validating the mechanisms regulating myosin II function and in studies on tumor migration and invasiveness. For this goal, several tools that allow for myosin II activation have been developed^24,27,66,67^. Many of the recent methods that allow acutely inducible activation of myosin II rely on optogenetic or optochemical tools. Despite allowing for precise spatiotemporal control, most of these tools act upstream of Rok (i.e., light-controlled activation of small GTPase RhoA or the optochemical control of Ca^2+^ levels), exhibiting more pleiotropic effect on cell contractility^96–100^, and lacking the specificity we present here. Also, we show that the destabilized variant of the engineered N-Rock (N-Rok::dGBP1) allows for minimal interference in *in vivo* studies when the target fusion protein is not present in the cell. Using destabilized nanobodies as fusion partners to reduce non-specific activities is a major step closer to using ever more specific tools to manipulate protein function.

### Effect on tracheal cell intercalation

Past work from our lab has shown that tracheal cell intercalation relays on the mechanical pulling of the stalk cells by leading tip cells and does not require actomyosin contractility ^44^. During the extensive junctional rearrangements, cells establish autocellular junctions, which are cell self-contacts. Somewhat surprisingly, previous studies have shown that autocellular contacts are normally eliminated in cultured cells through cell-self fusion which requires actomyosin contractility^101^. Also, exclusion of E-cadherin from contact sites has been proposed to be involved in eliminating self-contacts in different cultured cells^101,102^. We therefore hypothesized that the persistence of cell self-contacts in tracheal branches might result from insufficient levels of active actomyosin, thereby preserving the autocellular junctions.

Interestingly, herein we showed that ectopic activation of actomyosin with engineered Rok kinase did not prevent cell intercalation nor did it interfere with the formation of autocellular junctions. Nevertheless, we observed a gradation in the observed phenotypes ranging from normal branch elongation and cell intercalation (despite evident ectopic points of myosin activation at cell-cell contract), to more severe phenotypes displaying aberrant contacts between the tip cells or even detachment of some of the migrating cells from the rest of the branch resulting in aberrant branch network. Strikingly, we only observed an increase in pSqh along intercellular junction and not along autocellular junctions upon the expression of synthetic Rok. It thus remains possible that autocellular junctions are maintained by preventing the accumulation of actomyosin particularly at these junctions, and this exclusion cannot be overcome with the synthetic kinase. The importance of precisely dissecting the role of actin and myosin II in junctional rearrangements and autocellular junction formation is emphasized by the recent findings that autocellular AJs are also present in the retinal and adult mouse brain microvasculature (^103^, Maarja Mäe and Christer Betsholt, unpublished data).

Of note, a recent study from the Llimargas lab has shown that moderate levels of constitutively active Diaphanous (Dia, a protein that plays important role in the nucleation and elongation of F-actin filaments) disturbed cell elongation but not cell intercalation, while high levels of constitutively activated Dia impaired both cell elongation and intercalation, resulting in altered overall tracheal morphology^104^. Their data and ours (this manuscript) support the notion that cortical tension gradients resulting from local modulation of different pools of actomyosin lead to various impact on cell fitness/deformation. Further work is needed to fully understand the effect of modulating different pools of actin on cell intercalation and elongation in tracheal branches.

### Limitations of the study

Our results with engineered N-Rok in *Sqh::GFP* embryos showed that active N-Rok variants phosphorylate its target as early as expression starts (prominent Sqh::GFP foci are already observed at stage 11/12 when the *btlGal4* driver becomes active). Therefore, the driver used determines the spatial and temporal window of the activity of the synthetic kinase. An important point to be considered when employing the binder-based engineered kinases is the kinetics and reversibility of the phosphorylation by endogenous phosphatases influencing the total amounts of phosphorylated target protein in the tissue of choice at a given time.

It has been postulated that the range of possible targets is restricted by the structure of the kinase domain itself^105^. Although in principle our paradigm should be relatively easily extrapolated for use with different kinases and in other model systems, adding a protein binder to a catalytic domain of a kinase may influence protein folding, resulting in a perturbation in the positioning of the target with respect to the catalytic site and thus preventing phosphorylation. Hence, our system should be carefully tested for each kinase-substrate pair separately, and some fine-tuning (i.e., optimizing the linker length and amino-acid composition) might be needed for optimal tool performance.

### Future perspectives

Further possible avenues for expanding on the engineered kinases and other enzymes include phosphorylation of target proteins in specific subcellular compartments, for example only in the cell apical domain. During the course of this study, we have made use of a N-Rok synthetic kinase fused with an apical localization domain (*ALD::N-Rok::vhhGFP4-HA*^ZH-86Fb^) to restrict the activity of N-Rok to the apical cortex in interphasic *Drosophila* neuroblasts^106^. An important future improvement of protein binder-based synthetic biology would be to make target binding acutely inducible, either by light or chemically (add references, ask Alessandra). Combining the target-specific action presented in this manuscript with acute control over the activity of the fusion protein would allow an unprecedented level of spatiotemporal control of particular kinase-substrate interaction. Due to the large collection of available Drosophila strains with in-frame GFP or YFP insertions, our engineered Rok kinase (or, in broader context, any other chosen kinase-protein binder against GFP/YFP) can now be easily tested on other GFP fusion targets. Furthermore, carefully designed screens might identify other targets of protein kinases among the many GFP fusion lines available. Another intriguing avenue would be to define whether such engineered kinases can be reprogrammed to phosphorylate a specific target, which does not belong to the natural substrates of a given kinase, providing means for rewiring of the existing signaling pathways in cell or in the entire organism. Pioneering work on re-engineered PAK4 kinase^107^, DYRK1A kinase^108^ and Pim1 kinase^109^ provided proof-of-principle for re-programing kinase to recognize a novel substrate, in which the recognition site and motif is well known.

In summary, we present here the generation and the detailed characterization of a promising tool to control kinase-target protein activity and show a clear relationship between a specific kinase-target pair and cellular behavior in morphogenetic processes *in vivo*.

## Supporting information

Supplementary Movie 1_Dorsal closure

Supplementary Movie 4_Dorsal closure

Supplementary Movie 5_Dorsal closure

Supplementary Movie 7_Trachea development

Supplementary Movie 10_Intercalation_Rok_dGBP1

Supplementary Movie 9_Intercalation_control

Supplementary Movie 2_Dorsal closure

Supplementary Movie 3_Dorsal closure

Supplementary Movie 8_Branch elongation

Supplementary Movie 6_Dorsal closure

## Acknowledgements

We thank the Bloomington Stock Center, Giorgos Pyrowolakis, Stefan Luschnig and Robert Ward for providing fly stocks and antibodies; Bénédicte Sanson for the access to unpublished results; the Biozentrum Imaging Core Facility for maintenance of microscopes and support. The work in the laboratory of M.A. was supported by grants from the Swiss National Science Foundation (310030_192659/1) and by funds from the Kantons Basel-Stadt and Basel-Land.

## Supplementary Material

**Supplementary Data 1:** Multi FASTA file corresponding to all the inserts used in this work, and cloned in pUAST or pUASTattB. By virtue of the rich text encoding of this file (.rtf), the FASTA format is augmented with colored fonts so that the different domains the inserts contain are easily located; bold characters = kinase domains; green (bluish green 0,158,115) or red (vermillion 213,94,0) background = fluorescent protein binders; underlined characters = apical localization domain of Inscuteable; black background = Human influenza hemagglutinin tag; purple (reddish purple 204,121,167) background = mCherry. The restriction sites used for cloning are indicated with purple (reddish purple 204,121,167) characters.

## Supplementary Figure legends

**Supplementary Fig. 1**

All panels show stills from live-imaging with dorsal views of either *sqh_Sqh::GFP* or *sqh_Sqh::mCherry* embryos at stage 13/14 (dorsal closure) expressing either N-RokDead variants or vhhGFP4 nanobody alone expressed in the *en* domain (visualized by co-expression of mCherry-nls). Note the enhancement of fluorescent signal due to the binding of vhhGFP4 or 2m22 to GFP and to mCherry, respectively. The right panel shows magnification of the respective images for each of the embryo genotypes shown on the left. Please note, that the “control” image is the same as on Figure 3a and 4b, to have a single reference image for all N-Rok variants used. For every expressed synthetic kinase, the number of considered embryos is indicated (n). Scale bars: 50 μm, in the zoomed panel 10 μm..

**Supplementary Fig. 2**

All panels show stills from live-imaging with dorsal views of the developing *sqh_Sqh::GFP* or *sqh_Sqh::mCherry* embryos at stages 13/14-16 (dorsal closure) expressing variants of the synthetic kinase or the vhhGFP4 binder alone in the *en* domain (visualized by co-expression of mCherry-nls). Bottom panel shows dorsal views of embryo expressing Sqh^E20E21^ under the control of *enGal4* driver. Yellow arrows pointing to the Sqh::mCherry foci and actomyosin cable invaginations in N-Rok::2m22 panel. Please note, that the “control” image is the same as on Figure 3a, 4b and Supplementary Fig 1, to have a single reference image for all N-Rok variants used. For every genotype, the number of considered embryos is indicated (n). Scale bars: 50 μm.

**Supplementary Fig. 3**

N-Rok::vhhGFP4^Vi^ expression induces a partially missing cross vein phenotype (arrows). Control wing (left panel) compared to a wing in which N-Rok::vhhGFP4^Vi^ is expressed in the posterior compartment (right panel). The proximal direction (PR), the anterior and posterior cross veins (acv and pcv), as well as the anterior-posterior boundary (red dashed line) are indicated in the left panel. The wings shown come from female flies. Scale bar: 1 mm.

**Supplementary Fig. 4**

Panels show lateral views of fixed *sqh_Sqh::GFP* (top panel) or *sqh_Sqh::mCherry* (bottom panel) embryos at stage 13-14 (dorsal closure) expressing N-Rok::2m22 in the *en* domain (visualized by coexpression of mCherry-nls in the top panel). Embryos were stained with anti-phospho-myosin antibody. The panel on the right shows magnification of the respective Sqh and p-Sqh images for the presented embryos. Yellow arrows point to Sqh::mCherry and p-Sqh foci. The number of embryos analysed is indicated (n). Scale bars: 20 μm, in the zoomed panel 10 μm.

**Supplementary Fig. 5**

Expression of N-Rok::vhhGFP4^Vi^ in *Tkv::YFP* embryos has no effect on pMad levels. a) Panels show lateral views of fixed *Tkv::YFP* embryos at stage 13/14 (dorsal closure) expressing N-Rok::vhhGFP4^Vi^ in the *en* domain (visualized by co-expression of mCherry-nls). Embryos were stained with anti-phospho-Mad antibody. b) Panels show lateral views of fixed embryos at stage 13/14 (dorsal closure) expressing Tkv::GFP and N-Rok::vhhGFP4 in the *en* domain. Embryos were stained with anti-phospho-Sqh antibody. c) Panels show stills from live-imaging with dorsal views of developing *cadherin::GFP* embryos at stages 13/14-16 (dorsal closure) expressing N-Rok::vhhGFP4^Vi^ in the *en* domain (visualized by co-expression of mCherry-nls). d) Panels show lateral views of fixed *cadherin::GFP* embryo at stage 13/14. Embryos were stained with anti-phospho-myosin antibody. Note the enhancement of fluorescent signal due to the binding of vhhGFP4 to *cadherin::GFP* in the *en* domain. The number of considered embryos is indicated (n). Scale bars: 20 μm in a, b, d; 50 μm in c.

**Supplementary Table 1.** Viability and/or adult fly phenotypes induced by the expression of different synthetic kinase constructs with several Gal4 lines in the absence of the GFP-target. For every non-lethal condition, more than 20 adult flies were generated with consistent phenotypes. The genomic insertions and the Gal4 drivers were respectively sorted from left to right and top to bottom in descending strength of the phenotypes they induced. See text for details and Supplementary Fig. 3 for an illustration of the partially missing crossvein wing phenotype.

|  | Genomic insertion |  |  |  |  |
| --- | --- | --- | --- | --- | --- |
| Gal4 driver | *Rok-cat<sup>T2A</sup>* | *N-Rok::vhhGFP4<sup>ZH-86Fb</sup>* | *Rok-CAT<sup>3.1</sup>* | *N-Rok::vhhGFP4<sup>Vi</sup>* | *N-RokDead::vhhGFP4<sup>ZH-86Fb</sup>* |
| *tubGal4* | lethal | lethal | lethal | lethal | wild type |
| *enGal4* | lethal | lethal | missing crossveins | partially missing crossveins | wild type |
| *daGal4* | lethal | lethal | wild type | wild type | wild type |
| *69BGal4* | lethal | semi lethal | missing crossveins | wild type | wild type |
| *elavGal4* | rough eyes | rough eyes | wild type | wild type | wild type |

