## Supplementary Figures for "*In vivo* regulation of fluorescent fusion proteins by engineered kinases"

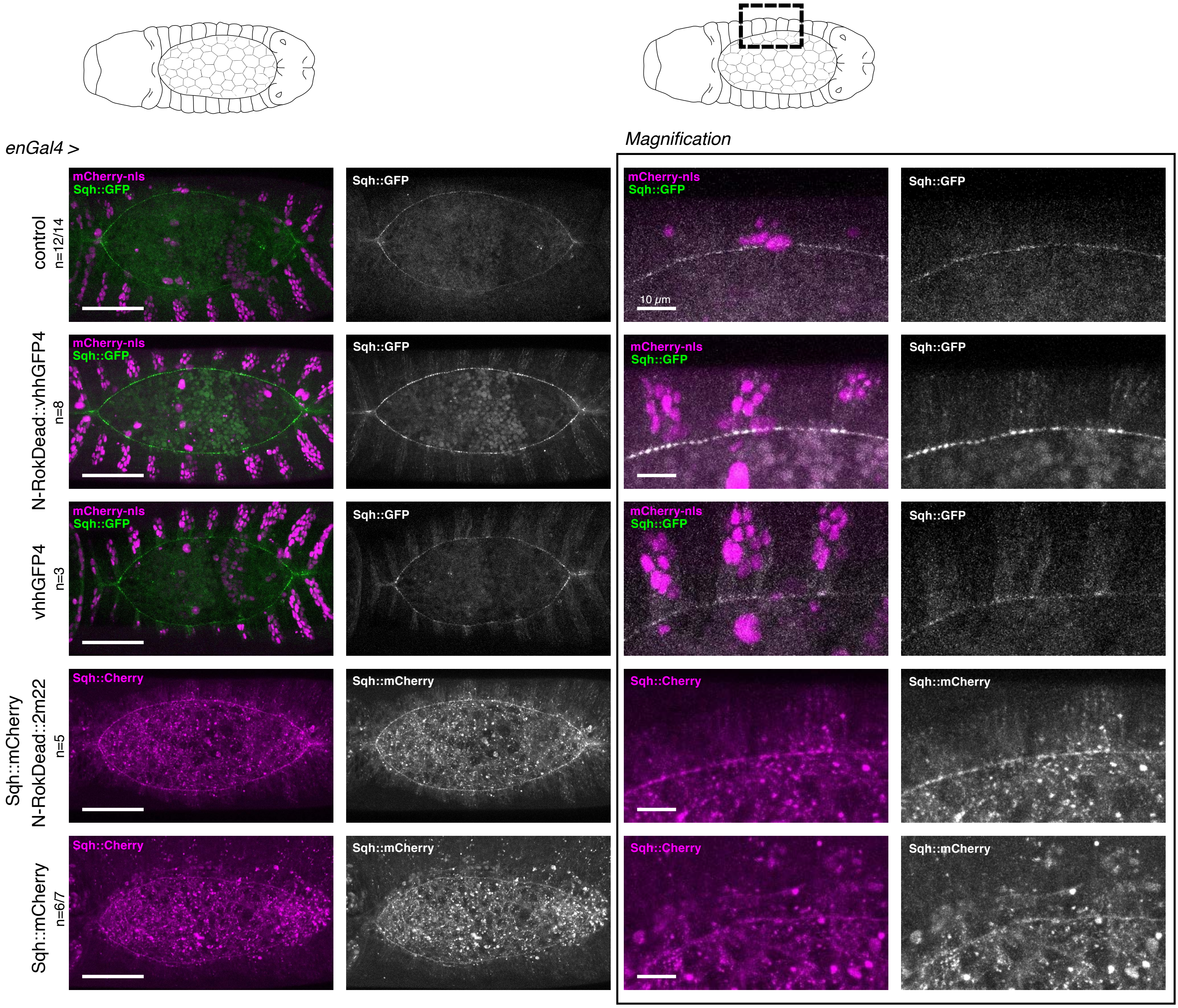


Supplementary Figure 1

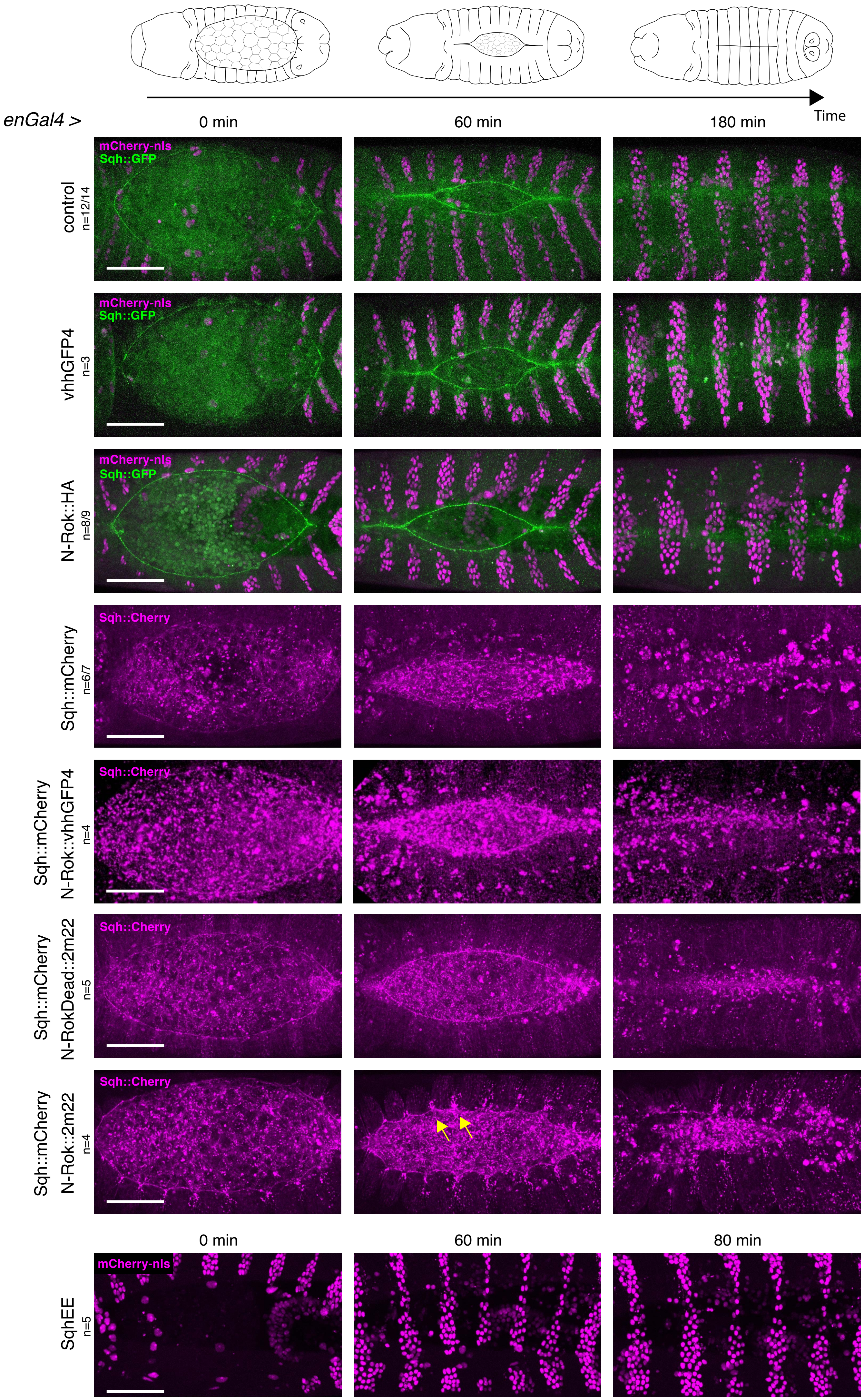


Supplementary Figure 2

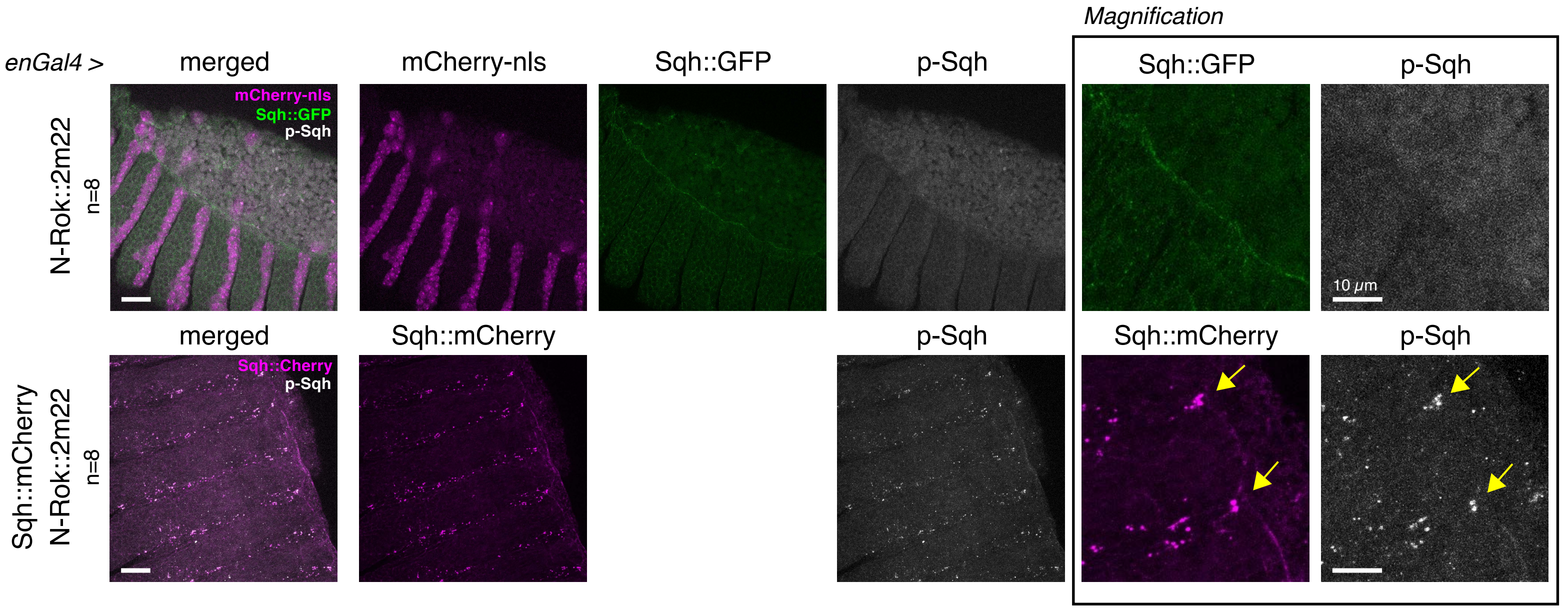


Supplementary Figure 4

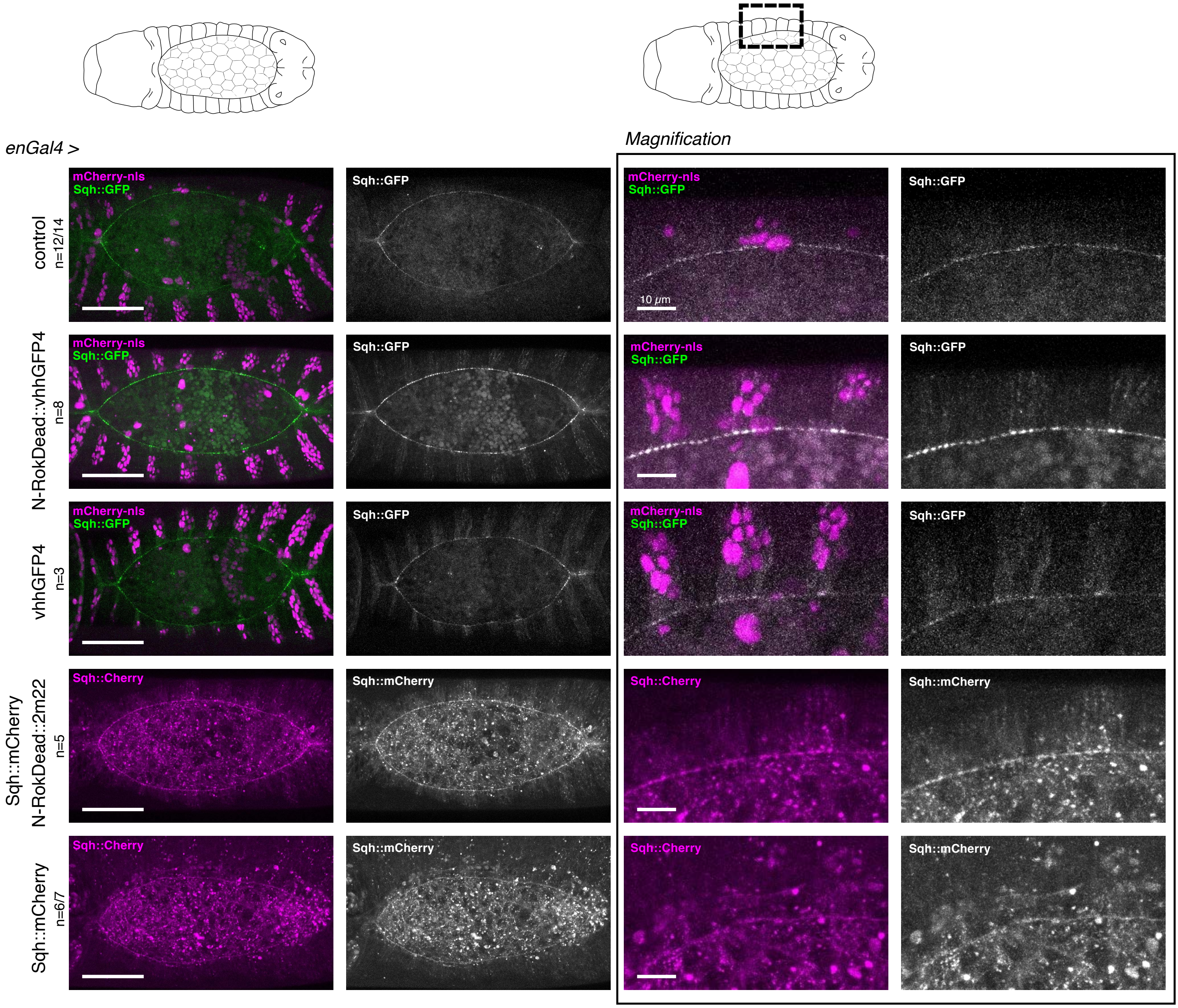


Supplementary Figure 1

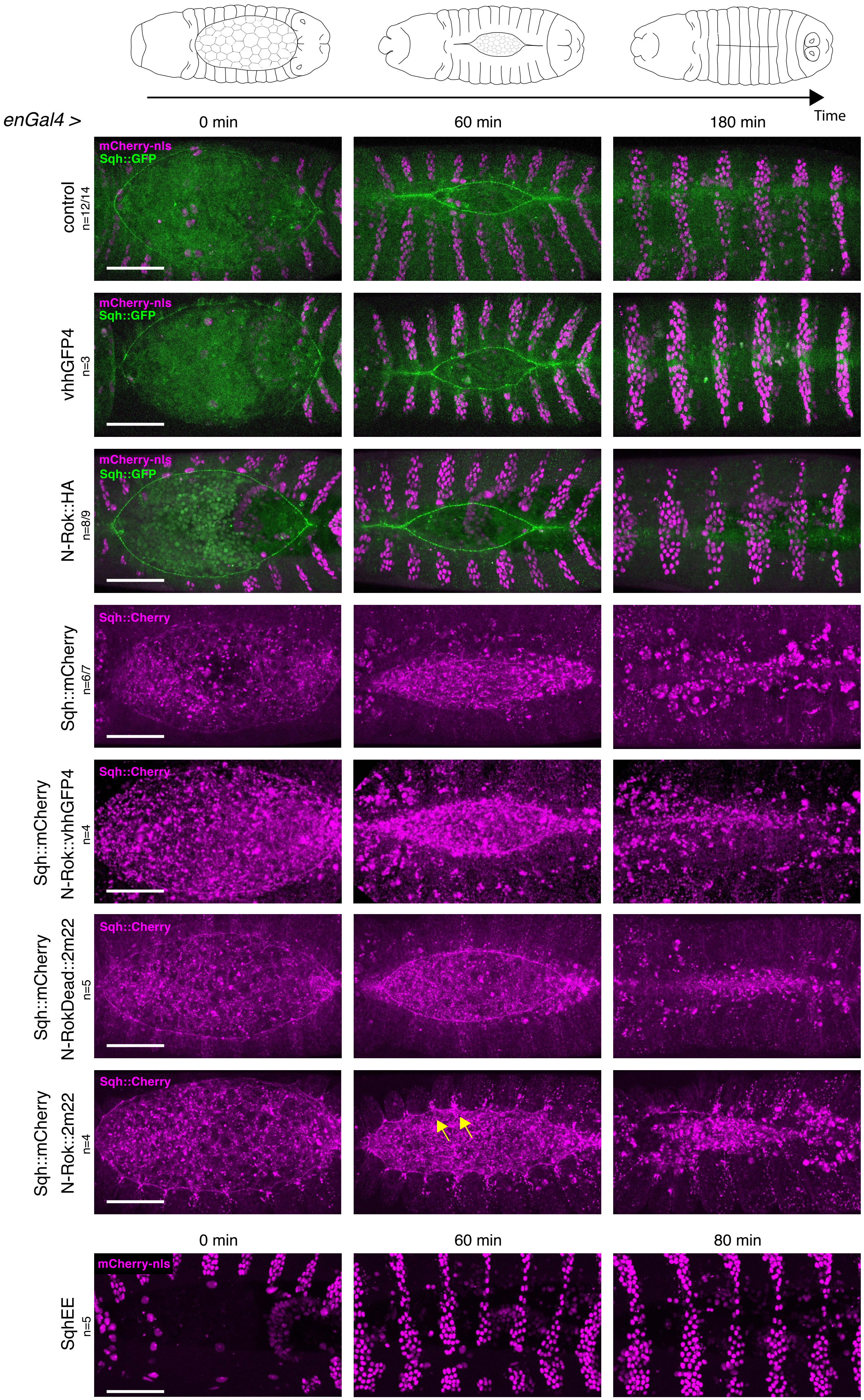


Supplementary Figure 2

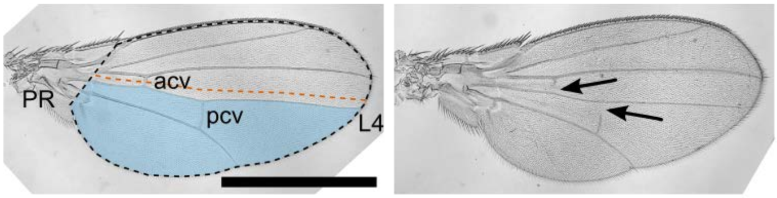


*yw*

*enGal4, N-Rok::vhhGFP4[Vi] / +*

Supplementary Figure 3

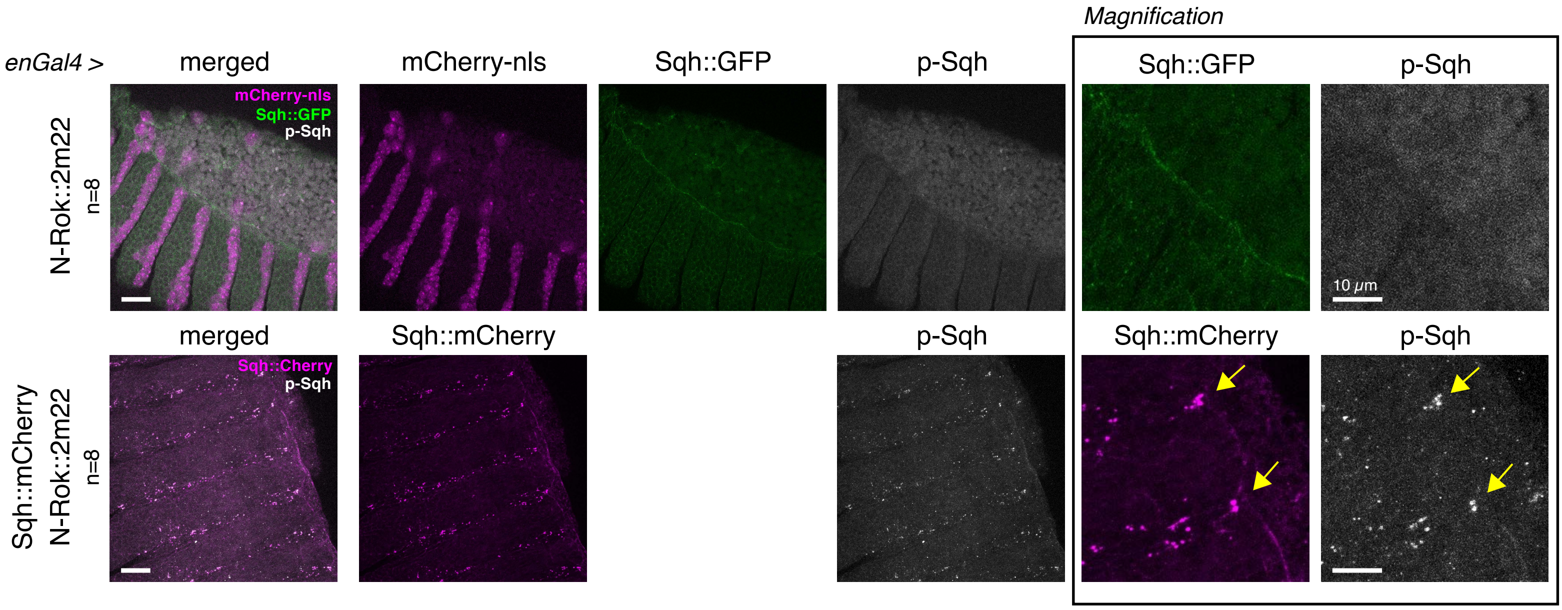


Supplementary Figure 4

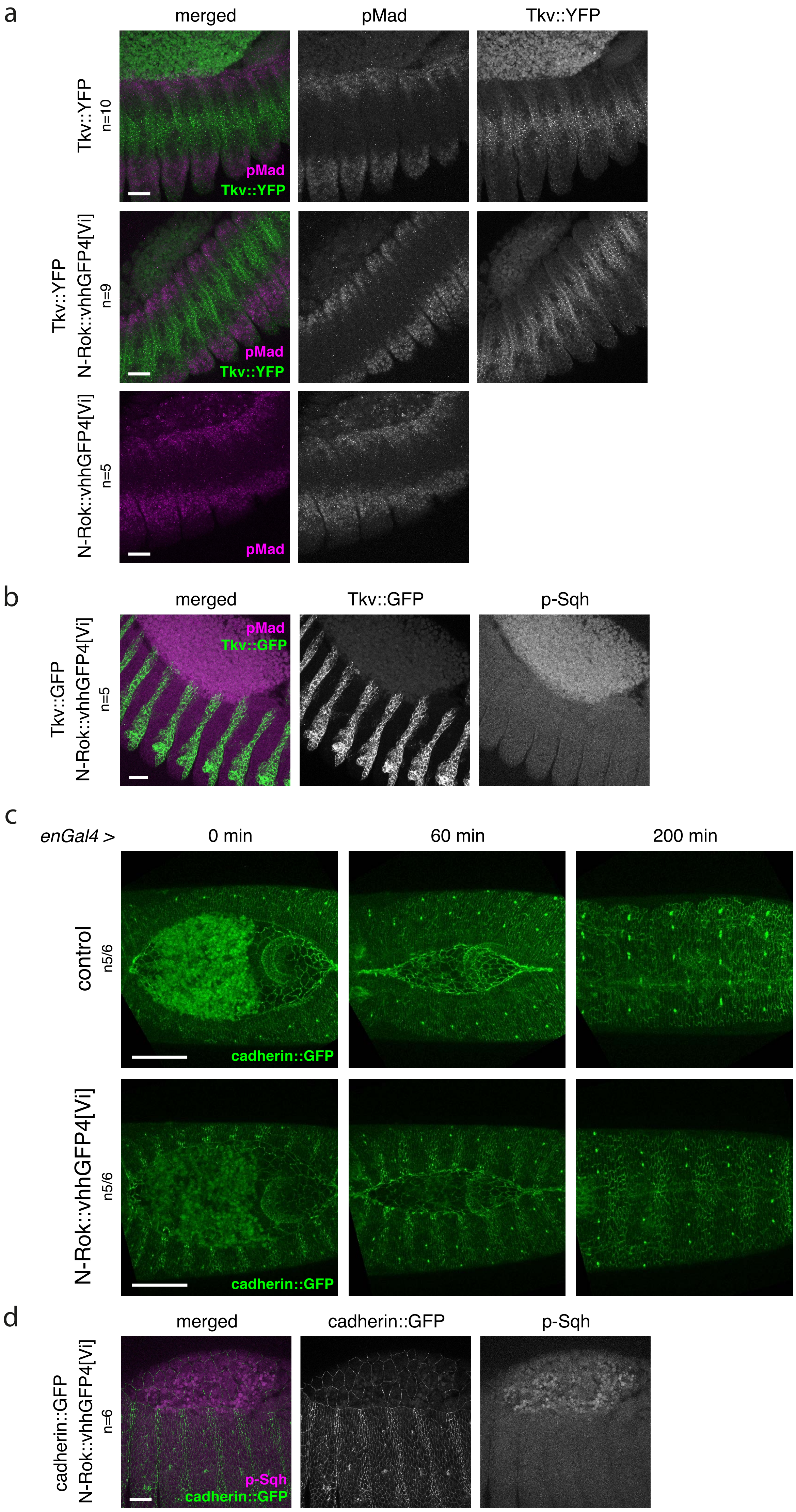
a
merged
pMad
Tkv::YFP
Tkv::YFP
n=10
pMad
Tkv::YFP
Tkv::YFP
N-Rok::vhhGFP4[Vi]
n=9
pMad
Tkv::YFP
N-Rok::vhhGFP4[Vi]
n=5
pMad
b
merged
Tkv::GFP
p-Sqh
Tkv::GFP
N-Rok::vhhGFP4[Vi]
n=5
pMad
Tkv::GFP
c
enGal4 >
0 min
60 min
200 min
control
n5/6
cadherin::GFP
N-Rok::vhhGFP4[Vi]
n5/6
cadherin::GFP
d
merged
cadherin::GFP
p-Sqh
cadherin::GFP
N-Rok::vhhGFP4[Vi]
n=6
p-Sqh
cadherin::GFP
